# c-Myb expression is critical to maintain proliferation and glucose metabolism of large pre-B cells

**DOI:** 10.1101/2020.09.09.290346

**Authors:** Andrea R. Daamen, Rowena B. Crittenden, Timothy P. Bender

## Abstract

The c-Myb transcription factor is required for the differentiation of CD19^+^ B-lineage cells and plays significant roles from the specification of the B cell lineage to the survival of pro-B cells. c-Myb coordinates the survival of pro-B cells with the expression of genes required for transition to the large pre-B cell stage of differentiation. However, it is not known if c-Myb is important for the proliferative expansion or subsequent differentiation into small pre-B cells. Here we demonstrate that c-Myb expression is important for large pre-B cell survival, proliferation, and differentiation into small pre-B cells. Utilizing genome-wide analysis, we found that c-Myb was important for maintaining glucose uptake and utilization and exogenous expression of Glut1 and Hk1 rescued large pre-B cell recovery and survival. Furthermore, we found that c-Myb is important for repression of Ikaros and Aiolos and our c-Myb-dependent gene signature was enriched in an Ikaros footprint of genes that drive cell cycle exit and the large to small pre-B cell transition. However, upon loss of c-Myb expression, inhibition of Ikaros activity was able to restore certain Ikaros-mediated gene expression changes but was insufficient to rescue recovery of large pre-B cell numbers. We found that c-Myb regulates glucose utilization and glucose-dependent survival through Hk1 in an Ikaros-independent manner. Thus, c-Myb regulation of glucose metabolism is critical to maintain large pre-B cell survival while repression of the Ikaros-mediated gene expression program is critical to prevent premature cell cycle exit and premature differentiation into small pre-B cells.

## Introduction

B cell development begins in the bone marrow of adult mammals from hematopoietic stem cells (HSC), and proceeds through a series of developmental stages driven by a network of cytokines and transcription factors, during which they gain expression of B-lineage specific genes while conversely losing the potential for differentiation into other myeloid or lymphoid cell types (1). Upon commitment to the B-cell lineage at the CD19^+^ pro-B cell stage, progenitors face two major developmental checkpoints defined by products of V(D)J recombination events at the *IgH* and *IgL* loci. The first of these checkpoints, the pre-BCR checkpoint, selects pro-B cells based on successful rearrangement at the *Igh* locus, production of a μ-H chain, and expression on the cell surface with the surrogate L chain and signaling molecules Igα and Igβ to form the pre-BCR. Upon expression of the pre-BCR, pro-B cells differentiate to large pre-B cells, which undergo a transient proliferative burst, exit the cell cycle, and differentiate into quiescent, small pre-B cells (2, 3). The second developmental checkpoint selects small pre-B cells for successful completion of VJ recombination at the *Igκ* L chain locus to produce a L chain that pairs with the μ-H chain and forms membrane IgM on the surface of immature B cells.

The successive differentiation steps from pro-B to large pre-B to small pre-B cells involves changes in proliferative and metabolic status, in particular changes in their reliance on glucose uptake and metabolism (4, 5). Pro-B cells maintain homeostatic levels of proliferation and metabolism while alternating between dividing populations engaged with IL-7-producing stromal cells and non-dividing populations that carry out V(D)J rearrangement at the IgH locus (2, 6, 7). In pro-B cells, glucose uptake is primarily utilized through oxidative phosphorylation over glycolysis (5). Subsequently, with expression of the pre-BCR, large pre-B cells undergo a limited burst of proliferation fueled by increased glucose uptake and increased overall metabolic activity. Similar to other highly proliferative cell types, large pre-B cells shift the majority of glucose utilization to glycolysis and the production of macromolecules to facilitate increased proliferation (5, 8). Then, after a limited number of cell divisions, large pre-B cells exit the cell cycle and repress glucose metabolism in preparation for differentiation into quiescent small pre-B cells. Repression of proliferation and metabolism is critical to minimize the risk of possible mutations that can occur during the process of VJ recombination in small pre-B cells. Similar to pro-B cells, small pre-B cells also rely on oxidative phosphorylation for energy production, however, overall levels of glucose uptake and metabolic activity are lower than in pro-B cells or large pre-B cells (4).

Signaling through the IL-7R and pre-BCR are the major drivers of development across the pre-BCR checkpoint including changes in gene expression and shifts in proliferative and metabolic status (2, 3). Pro-B cells mainly rely on IL-7R signaling through the Jak/Stat and Plcγ/DAG/Pkc pathways for cell survival, but also to maintain cell proliferation and metabolism (9–12). Stat5 mediates pro-B cell survival through regulating the balance between pro-and anti-apoptotic Bcl-2 family members and contributes to proliferation through induction of cyclin D3 expression (9, 13). However, the majority of cyclin D3 expressed in pro-B cells is not involved in proliferation and *Ccnd3*^*-/-*^ pro-B cells only exhibit a moderate reduction in >2n DNA content (14, 15). Plcγ signals through the second messenger DAG to activate Pkc and ultimately mTORC1 and this pathway is critical for pro-B cell proliferation and metabolism. Upon its expression on large pre-B cells, the pre-BCR initially synergizes with the IL-7R to increase sensitivity to IL-7 (16, 17). Activation of Jak/Stat5 signaling downstream of the IL-7R in combination with PI3K/Akt signaling downstream of the IL-7R and pre-BCR lead to the expression of cyclin D3, expression of c-Myc, and activation of mTORC1 to promote enhanced proliferation and metabolism (15, 18, 19). Subsequently, a threshold level of pre-BCR signaling shifts large pre-B cells from a state of proliferation to differentiation. In particular, pre-BCR-mediated induction and activation of Blnk (SLP-65) inhibits PI3K/Akt proliferative signaling and induces Irf4, which, in turn, induces expression of the Ikaros transcription factor family members Ikaros (*Ikzf1*) and Aiolos (*Ikzf3*) (20). Ikaros family members form homo-or heterodimers to facilitate optimal DNA binding and activity and, together, Ikaros and Aiolos mediate the majority of gene expression changes that occur at the large to small pre-B cell transition (21, 22).

The c-Myb transcription factor (*Myb*) is a DNA-binding protein that acts as both a transcriptional activator and repressor (23, 24). c-Myb is abundantly expressed by HSC and immature progenitors of each hematopoietic lineage and expression is decreased or ablated in mature hematopoietic cells (25). *Myb* null mutations are embryonic lethal by day 15 post-coitus due to severe anemia caused by the failure to undergo adult erythropoiesis in the fetal liver (26). To overcome the embryonic lethality of *Myb*-null mutations, Cre/LoxP based conditional deletion has been used to demonstrate that c-Myb is important for multiple stages of B-cell development (27–29). Ablation of c-Myb expression in early B-lineage progenitors using *Myb*^*f/f*^ *Mb1-cre* mice demonstrated that c-Myb is absolutely required for the development of CD19^+^ B-lineage cells and that c-Myb is crucial for the proper expression of IL-7Rα and Ebf1 (28, 29). Loss of c-Myb expression in *Myb*^*f/f*^ *CD19-cre* mice, where Cre-mediated deletion occurs in late pro-B cells, results in a partial block to differentiation beyond the pro-B cell stage and a decreased number of large and small pre-B cells as well as immature B cells (27). We recently reported that c-Myb is required for the survival of CD19^+^ pro-B cells where c-Myb sets baseline expression of the proapoptotic proteins Bim and Bmf and is further required for expression of IL-7Rα, which suppresses expression of Bim and Bmf, thus regulating the lifespan of pro-B cells (30). However, rescue of c-Myb-deficient pro-B cell survival with a Bcl-2 transgene does not rescue the large and small pre-B cell compartments. c-Myb also controls the expression of genes that are required for the pre-BCR checkpoint including components of the surrogate light chain and cyclin D3 (30). However, it is not known if c-Myb is important in the large pre-B cell compartment or differentiation to the small pre-B cell compartment.

We now demonstrate that c-Myb plays a distinct role in the large pre-B cell compartment and found that loss of c-Myb expression resulted in decreased large pre-B cell proliferation and survival. Genome-wide analysis of c-Myb function by RNA-seq revealed enrichment of c-Myb-regulated genes in pathways related to metabolism and glucose utilization. Furthermore, exogenous expression of proximal components of glucose utilization, Glut and Hk1, restored recovery and decreased apoptosis of c-Myb-deficient large pre-B cells. We also identified a role for c-Myb in repression of Ikaros and Aiolos and demonstrate that the c-Myb-dependent transcriptional profile was highly enriched in Ikaros targets that are important for mediating cell cycle exit and the large to small pre-B cell transition. However, while expression of a dominant negative Ikaros mutant was able to reverse gene expression changes induced by c-Myb knockdown and the c-Myb-dependent increase in Ikaros expression, it was insufficient to fully rescue recovery of c-Myb deficient large pre-B cells, suggesting that increased Ikaros expression cannot account for all aspects of the c-Myb-dependent large pre-B cell phenotype. We found that c-Myb regulated Hk1 mRNA expression, glucose utilization and glucose-dependent survival, in an Ikaros-independent manner. Thus, c-Myb exhibits both Ikaros-dependent and Ikaros-independent functions in large pre-B cells in order to maintain large pre-B cell metabolism and proliferation while preventing the premature initiation of gene expression changes that promote differentiation.

## Materials and Methods

### Mice

*Myb*^*ff*^ and *Bcl2Tg* mice have been previously described (31–33). *Rag2*^*-/-*^ (Taconic Farms), *Myb*^*ff*^ *Rag2*^*-/-*^, and *Myb*^*ff*^ *Rag2*^*-/-*^ *Bcl2Tg* mice were bred at the University of Virginia. Mice were housed at the University of Virginia in a barrier facility and were 6-8 weeks old when used for experiments. These studies were approved by the Institutional Animal Care and Use Committee (IACUC) at the University of Virginia.

### Retrovirus Vectors

The retrovirus vectors, pMIG-R1, pMSCV-IRES-tNGFR, pMSCV-IRES-tNGFR-Cre, pMIG-17.2.25, pSIREN-RetroQ-shLuc-IRES-eGFP, pSIREN-RetroQ-shMyb-IRES-eGFP, pMIG-IkWT, pMIG-Ik159A, pMSCV-IRES-tNGFR-Glut1, and pMSCV-IRES-tNGFR-Hk1 have been previously described (22, 34–39). pMIG-IkWT and pMIG-Ik159A were provided by Dr. Hilde Schjerven (University of California, San Francisco). pMSCV-IRES-tNGFR-Glut1 and pMSCV-IRES-tNGFR-Hk1 were provided by Dr. J. Rathmell (Vanderbilt University). To produce pSIREN-RetroQ-shMyb-IRES-tNGFR, the IRES-tNGFR from pMSCV-IRES-tNGFR was cloned into the BglII/XhoI site of pLITMUS28, then into the EcoRV site of pSIREN-RetroQ-shMyb. Retrovirus supernatants were generated by transient transfection of HEK-293T cells and titered on NIH-3T3 cells as described previously (34).

### Cell Culture and Retrovirus Transduction

For de-novo generation of pre-B cells from *Myb*^*ff*^ *Rag2*^*-/-*^ and *Myb*^*ff*^ *Rag2*^*-/-*^ *Bcl2Tg* mice, pro-B cells were positively selected from bone marrow using anti-CD19-labeled magnetic beads (Miltenyi Biotec) and were cultured for 24h hr in Opti-MEM supplemented with 15% FBS (Life Technologies), 100 U/ml penicillin-streptomycin, 2mM L-glutamine, 50 μM 2-ME, and 5 ng/ml IL-7 (PeproTech). Cells were then transduced with pMIG-R1 or pMIG-17.2.25, returned to culture for 24 hr, transduced with pMSCV-IRES-tNGFR or pMSCV-IRES-tNGFR-Cre, returned to culture in Opti-MEM supplemented with 15% FBS (Life Technologies), 100 U/ml penicillin-streptomycin, 2mM L-glutamine, 50 μM 2-ME, and 10 ng/ml IL-7 (PeproTech), and analyzed 24, 48, and 72 hr later by flow cytometry.

*Irf4/8*^*-/-*^ large pre-B cells (M. Mandal, University of Chicago) were cultured on an OP9 stromal cell layer in Opti-MEM supplemented with 10% FBS (Life Technologies), 100 U/ml penicillin-streptomycin, 50 μM 2-ME, and 10 ng/ml IL-7 (PeproTech). The OP9 stromal cell line (D. Allman, University of Pennsylvania) was grown in DMEM supplemented with 10% FBS (Life Technologies), 100 U/ml penicillin-streptomycin, 2mM L-glutamine, 1% Sodium pyruvate, 1% 100X HEPES, 0.1% gentamycin, and 50 μM 2-ME. For transduction, *Irf4/8*^*-/-*^ large pre-B cells were seeded on OP9 cells in a 24-well plate, transduced or co-transduced with retroviral vectors, re-plated on OP9 cells in a 24-well or 96-well plate, and analyzed 24, 48, 72, 96, and 120 hr later by flow cytometry.

### Flow Cytometry and Cell Sorting

Pro-B cells were positively selected from bone marrow and transduced with pMIG-R1 or pMIG-17.2.25, returned to culture for 24 hr, then transduced with pMSCV-IRES-tNGFR or pMSCV-IRES-tNGFR-Cre. Co-transduced pro-B cells or *Irf4/8*^*-/-*^ large pre-B cells were maintained in culture, and 1-2 × 10^6^ cells were stained with fluorochrome-conjugated antibodies as previously described (27). Cells were analyzed on FACSCanto II (BD Immunocytometry Systems), Cytoflex (Beckman Coulter Life Sciences), or Attune NxT (Thermo Fisher Scientific) instruments. Total cells were determined using AccuCount Blank Particles (Spherotech). Flow cytometric data were analyzed using FlowJo software (TreeStar). Cell sorting was performed on FACSVantage SE Turbo Sorter or BD Influx Cell Sorter (BD Immunocytometry Systems). Antibodies and reagents were purchased from: eBioscience: 7-aminoactinomycin D (7AAD); BD Pharmingen: streptavidin-PE; Biolegend: Zombie Aqua, anti-CD271-APC (NGFR; ME20.4); Miltenyi: anti-LNGFR-Biotin. For anti-active Caspase 3 staining, cells were fixed and permeabilized using the BD Cytofix/Cytoperm Kit per the manufacturer’s protocol, then stained with Alexa Fluor 647 Rabbit Anti-Active Caspase-3 (Clone C92-605; BD Pharmingen) per manufacturer’s protocol. EdU staining was performed with a Click-iT Plus EdU Alexa Fluor 647 Flow Cytometry Assay Kit (Life Technologies) per the manufacturer’s protocol. Glucose uptake analysis was performed with a 2-NBDG Glucose Uptake Assay Kit (BioVision) per the manufacturer’s protocol. The glucose uptake inhibitor, phloretin, was utilized as a negative control.

### Hexokinase Activity Assay

*Irf4/8*^*-/-*^ large pre-B cells were transduced with pSIREN-RetroQ-shLuc-IRES-eGFP or pSIREN-RetroQ-shMyb-IRES-eGFP and electronically sorted based on the expression of GFP at 72 hr post-transduction. The enzymatic activity of hexokinase (HK) was measured using the Hexokinase enzyme assay kit (Biomedical Research Service and Clinical Application; E-111) per the manufacturer’s protocol.

### Quantitative Real-Time PCR

*Irf4/8*^*-/-*^ large pre-B cells were electronically sorted based on NGFR and/or GFP expression, total cellular RNA was homogenized using TRIzol Reagent (Invitrogen), and purified using the RNeasy Mini Kit (Qiagen) according to the manufacturer’s protocol. Treatment with RNase-free DNase I (Invitrogen) was used to remove contaminating genomic DNA and cDNA was prepared with the Superscript III First-Strand Synthesis System (Invitrogen). Quantitative RT-PCR (qRT-PCR) was performed on cDNA as previously described (30). Primers used are listed in Supplemental Table 1.

### RNA-seq

Retrovirus-transduced *Irf4/8*^*-/-*^ large pre-B cells were electronically sorted based on GFP expression, total cellular RNA was homogenized using TRIzol Reagent (Invitrogen), and purified using the RNeasy Mini Kit (Qiagen) according to the manufacturer’s protocol. RNA quantity was measured using a NanoView (GE Health Systems) and quality was assessed using the Agilent 4200 TapeStation RNA kit. mRNA was purified by NEBNext Poly(A) mRNA Magnetic Isolation Module (New England Biolabs) and converted into RNA-seq libraries using NEBNext Ultra II Directional RNA Library Prep Kit for Illumina (New England Biolabs) following the manufacturer’s protocol. Libraries were quantified with a Qubit 3 Fluorometer (Thermo Fisher Scientific) using a dsDNA HS kit (Thermo Fisher Scientific). Average library size was estimated by Agilent TapeStation. NGS was performed with the NextSeq 150 High Output sequencing kit reagent cartridge on the Illumina NextSeq 500 for paired-end sequencing. Quality assessment of the sequencing reads was performed by running FASTQC. The fastq files were aligned to the mouse genome mm10 followed by gene counting with HTSeq. Differential expression analysis was performed with the R/Bioconductor package DESeq2.

### Gene Set Analysis

Gene set enrichment analysis was performed utilizing the investigate function of the Molecular Signatures Database (MSigDb) (40–42). RNA-seq results at a threshold of p<0.0001 were used to compute overlaps with curated gene set collections for KEGG pathways and Gene Ontology (GO) gene sets for Biological Processes at an FDR q-value below 0.05.

### Statistical Analyses

Differences between data sets were analyzed using the two-tailed Student t test and at a confidence level of 95% for all experiments; error bars represent SEM. Data sets were analyzed and figures were prepared with Prism v.7.0 and Prism v.8.0 (GraphPad Software).

## Results

### c-Myb is critical for the proliferative expansion of large pre-B cells

We previously reported that c-Myb coordinates pro-B cell survival and proliferation with the expression of genes that are critical for initiation of the pre-BCR checkpoint (30). However, it is not known if c-Myb is important in the large pre-B cell compartment or transition to the small pre-B cell compartment. Due to the transient nature of the large pre-B cell compartment it is difficult to isolate this population for study *in vivo*. Thus, to examine c-Myb function in large pre-B cells, we have utilized two model systems. In the first model, *Myb*^*ff*^ *Rag2*^*-/-*^ pro-B cells were transduced in cell culture with a retrovirus (MIG-17.2.25) that encodes a rearranged mu heavy chain (μHC) and a GFP reporter as a bicistronic mRNA (μHC–GFP) to promote differentiation into large pre-B cells (36). Subsequently, these cells were transduced 24 h later with retroviruses encoding a tNGFR reporter or the tNGFR reporter and Cre (Cre-tNGFR) to ablate c-Myb expression (Fig. 1A). Transition to the large pre-B cell compartment was demonstrated by an increase in cell size and cell number over the time course of the experiment as compared to empty-vector-transduced pro-B cell controls (Supplemental Figures. 1A, 1B). Furthermore, by 48 h post transduction with µHC, a population of small pre-B cells appeared (Figure 2A). To determine if c-Myb is important in large pre-B cells, the relative recovery of MIG-µHC/tNGFR (µHC) and MIG-µHC/tNGFR-Cre (µHC/Cre) transduced pro-B cells was assessed over a 72 h time course (Fig. 1B). We found that the relative recovery of µHC/Cre transduced pro-B cells was decreased by approximately 80% compared to µHC transduced pro-B cells 48 h after transduction and >90% by 72 h post-transduction. Thus, c-Myb appears to be crucial for continued proliferative expansion of µHC-transduced pro-B cells.

**FIGURE 1.**
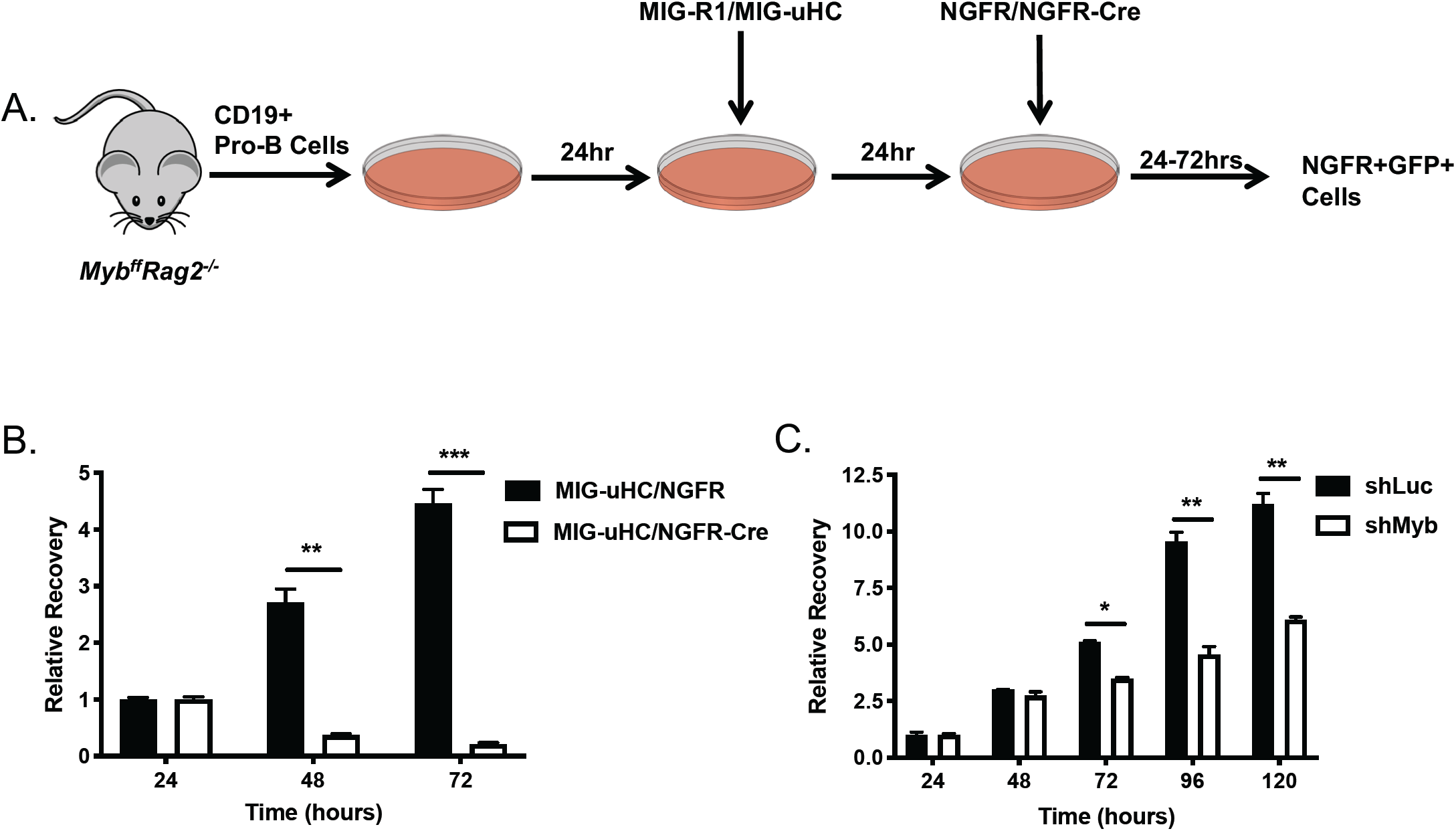
c-Myb Deletion in *Myb*^*ff*^*Rag2*^*-/-*^ *uHC*-Transduced Pro-B cells Leads to Decreased Pre-B Cell Survival and Proliferation. (**A**) Experimental model for in vitro generation of pre-B cells. CD19^+^ pro-B cells were positively selected on magnetic beads from *Myb*^*ff*^ *Rag2*^*-/-*^ mice and cultured for 24 h with 10ng/ml IL-7. These cells were subsequently transduced with MIG-17.2.25 (uHC) or the empty GFP reporter vector (MIG-R1) and 24 h later with NGFR-Cre or the empty NGFR reporter vector. Following retrovirus transduction, pre-B cells were cultured with 10ng/ml IL-7 and every 24 h, co-transduced GFP^+^ NGFR^+^ cells were analyzed for total cell numbers. To determine relative recovery, the total number of co-transduced cells was set as 1 and relative recovery was calculated as a ratio compared with the number of GFP^+^ NGFR^+^ cells at the 24 h time point. (**B**) Co-transduced GFP^+^ NGFR^+^ cells were analyzed at 24, 48, and 72 h post-transduction by flow cytometry and relative recovery was determined. Data are representative of 2 independent experiments. (**C**) *Irf4/8*^*-/-*^ large pre-B cells were transduced with shLuc-GFP or shMyb-GFP cultured with 10ng/ml IL-7. Total numbers of GFP^+^ cells was analyzed every 24 h post-transduction over a 120 h time course by flow cytometry. Relative recovery was determined as in (B). Data are representative of 3 independent experiments. All retrovirus transductions were performed with three replicates per condition. *p < 0.05, **p < 0.005, ***p < 0.0005.

**FIGURE 2.**
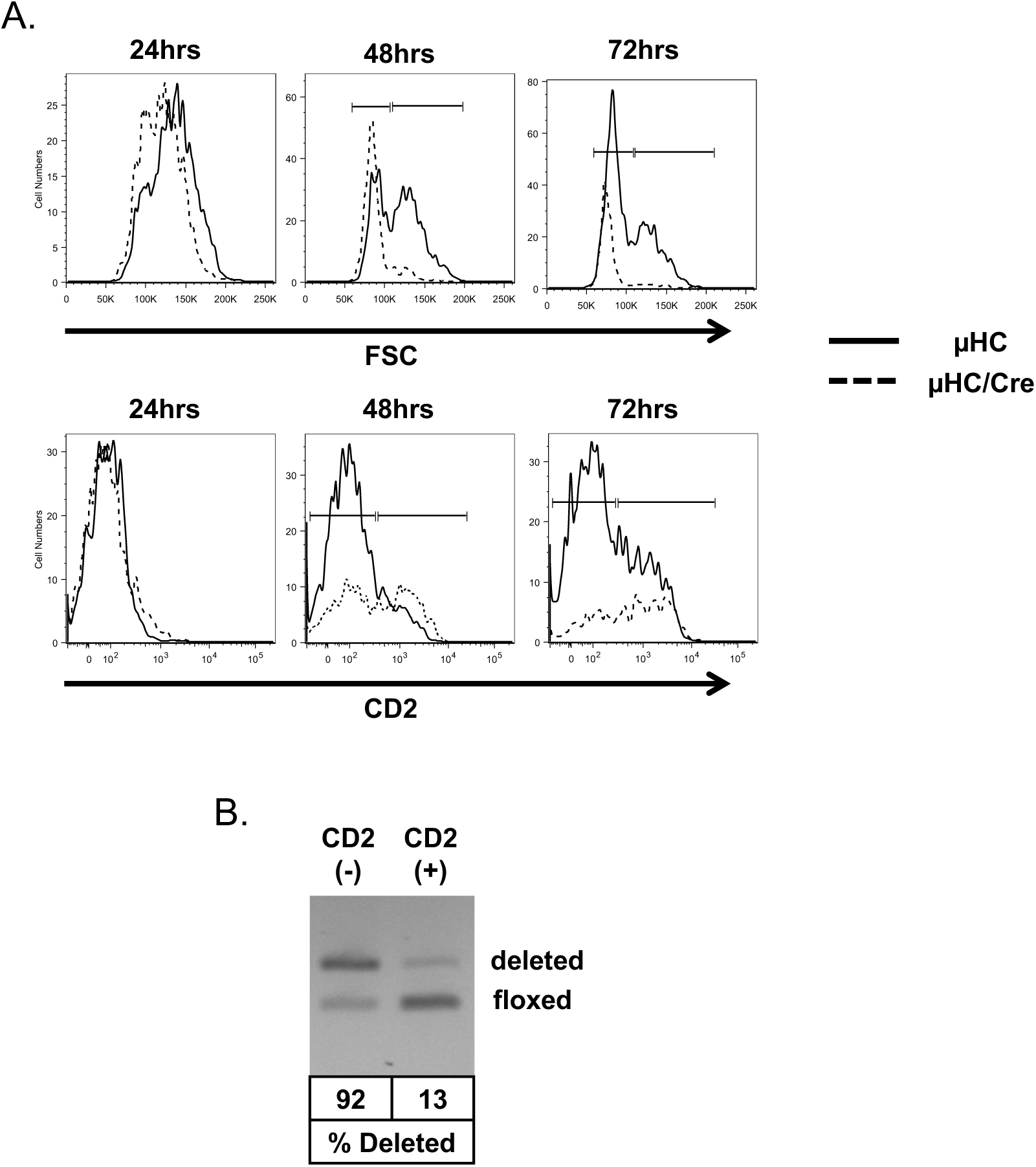
Defects in Proliferation and Selection against Differentiation of c-Myb-Deficient Pre-B Cells. Pre-B cells were generated by retrovirus transduction of *Myb*^*ff*^ *Rag2*^*-/-*^ pro-B cells as described in Fig. 1. (**A**) Representative histograms of FSC and CD2 expression of co-transduced GFP^+^ NGFR^+^ cells from the 48 h time point of the relative recovery time course in Figure 1B. Data are representative of 2 independent experiments. (**B**) GFP^+^ NGFR^+^ CD2^+^ cells were electronically sorted and deletion efficiency of the floxed *Myb* allele was analyzed by PCR as previously described (31). All retrovirus transductions were performed with three replicates per condition.

In this model, large pre-B cells continued to differentiate and by 48 h post-transduction with μHC, a population of small pre-B cells appeared (Fig. 2A). Since the μHC-transduction model ultimately produces a heterogeneous population of large and small pre-B cells, we also examined the *Irf4*^*-/-*^ *Irf8*^*-/-*^ double deficient (*Irf4/8*^*-/-*^) large pre-B cell line as a second model that represents large pre-B cells (43). This cell line is derived from *Irf4/Irf8*^-/-^ mice that accumulate cycling large pre-B cells in the bone marrow that fail to differentiate and undergo recombination at the IgL locus. However, as *Irf4/8*^*-/-*^ large pre-B cells are not on a *Myb*^*ff*^ background, they were transduced with retroviruses that co-produce a GFP reporter and a *Myb* mRNA targeting shRNA (shMyb) or a GFP reporter and a control *Luciferase* mRNA targeting shRNA (shLuc) (28, 38). Transduction of *Irf4/8*^*-/-*^ large pre-B cells with shMyb resulted in approximately an 80% knock-down of c-Myb protein (Supplemental Figure. 1C). In addition, shMyb transduction resulted in a 30% decrease in relative recovery at 72 h post-transduction. Due to the differences in phenotype between the shMyb-mediated knockdown and Cre-mediated knockout approaches, we extended the relative recovery time course from 72 h to 120 h in the shMyb-transduced *Irf4/8*^*-/-*^ large pre-B cell model. With these additional time points, we found a 50-60% decrease in recovery of shMyb-transduced *Irf4/8*^*-/-*^ large pre-B cells at 96 and 120 h post-transduction compared to shLuc transduced controls (Fig. 1C). Thus, the shMyb knockdown model in *Irf4/8*^*-/-*^ large pre-B cells behaves similar to a hypomorphic c-Myb mutant and is consistent with results obtained with c-Myb hypomorphic mutations that are viable but have a less drastic phenotype than *Myb* null mutations (44, 45). Taken together, these experiments demonstrate that c-Myb is important for the proliferative expansion of large pre-B cells.

### c-Myb is important for pre-B cell survival, proliferation and accumulation of CD2^+^ small pre-B cells

Large pre-B cells undergo a limited number of divisions, exit the cell cycle and differentiate into CD2^+^ small pre-B cells. The deficit in relative recovery that we observed in both the µHC-transduction and *Irf4/8*^*-/-*^ large pre-B cell models upon loss of c-Myb expression (Figs. 1B, 1C) could be due to increased apoptosis, lack of proliferation, premature differentiation from large to small pre-B cells or all three. To address these questions, we examined the ability of c-Myb deficient µHC-transduced pro-B cells to differentiate into small, CD2^+^ pre-B cells (Fig. 2A). *Myb*^*f/f*^ *Rag2*^*-/-*^ pro-B cells were transduced with MIG-µHC and 24 h later transduced with tNGFR (control) or tNGFR-Cre (Cre) retroviruses and cultured for an additional 24-72 h (Fig. 1A). At 24 h, the control µHC-transduced pro-B cells were mostly large pre-B cells, based on FSC, though a small proportion appeared to be small pre-B cells. By 48 and 72 h, the control µHC-transduced pro-B cell population increased the fraction of small pre-B cells. In contrast, the entire population of µHC/Cre transduced pro-B cells was smaller than the control cells 24 h post-transduction. By 48 and 72 h post-transduction the vast majority of µHC/Cre transduced pro-B cells were small. Furthermore, by 72 h post-transduction the µHC/Cre transduced cells displayed lower FSC than the control µHC-transduced pro-B cells. Thus, the µHC/Cre transduced pro-B cell population failed to maintain a large pre-B cell population. In addition, while control µHC transduced pro-B cells decreased in cell size, as measured by forward scatter (FSC), they also gradually acquired expression of the small pre-B cell marker CD2 over the 72 h time course. In contrast, the µHC/Cre transduced pro-B cell population contained a higher proportion of CD2^+^ cells at 48 and 72 h although the relative recovery of µHC/Cre transduced pro-B cells was much smaller at these time points (Fig. 1B), suggesting the possibility that the c-Myb deficient population quickly differentiated into small CD2^+^ pre-B cells. To determine if the floxed *Myb* locus was deleted in the CD2^+^ µHC/Cre-transduced pro-B cell population, we electronically sorted the CD2^+^ and CD2^-^ populations and measured the deletion efficiency at the *Myb*^*f/f*^ loci by PCR as we have previously (27, 31). Deletion at the *Myb*^*f/f*^ locus was very efficient (92%) in the CD2^-^ fraction (Fig. 2B). In contrast, deletion efficiency was poor (13%) in the CD2^+^ fraction, demonstrating a strong selection against loss of the floxed *Myb* allele in the CD2^+^ fraction, suggesting that c-Myb deficient large pre-B cells fail to transit to or survive in the CD2^+^ small pre-B cell fraction.

To better understand the nature of the defect in µHC-transduced pro-B cells, we measured apoptotic cell death by flow cytometry to detect active Caspase 3^+^ cells. Loss of c-Myb resulted in a 2 to 3-fold increase in the proportion of active Caspase 3^+^ µHC-transduced pro-B cells 24 and 48 h after tNGFR-Cre transduction (Fig. 3A). Similarly, c-Myb knockdown in *Irf4/8*^*-/-*^ large pre-B cells resulted in a 2-fold increase in active Caspase 3^+^ cells by 72 h and 96 h post-transduction with shMyb (Fig. 3B). These data indicated that ablation of c-Myb expression resulted in increased pre-B cell death. We also assessed DNA synthesis and DNA content by simultaneously measuring EdU uptake and 7AAD DNA staining. By 24 and 48 h post-transduction, we detected a >80% reduction in the proportion of S-phase µHC-transduced pro-B cells that were actively synthesizing new DNA compared to controls (Fig. 3C). This result demonstrated a severe defect in proliferation following c-Myb deletion. We also noted an increased number of c-Myb-deficient cells that by DNA content appeared to be in S-phase but did not label with EdU, possibly representing cells that have undergone an intra-S-phase cell cycle arrest, which typically occurs due to the activation of a DNA damage checkpoint (46). In the *Irf4/8*^*-/-*^ large pre-B cell model, we also detected an approximately 30% reduction in the proportion of S-phase cells that were actively synthesizing new DNA compared to controls at 96 h and 120 h after shMyb transduction (Fig. 3D), consistent with the less severe phenotype in this model. Thus, c-Myb is important for the survival and continued proliferation of large pre-B cells. Taken together, these results suggest that loss of c-Myb in large pre-B cells results in rapid loss of proliferation, cell death and a failure to transit to the CD2^+^ small pre-B cell compartment.

**FIGURE 3.**
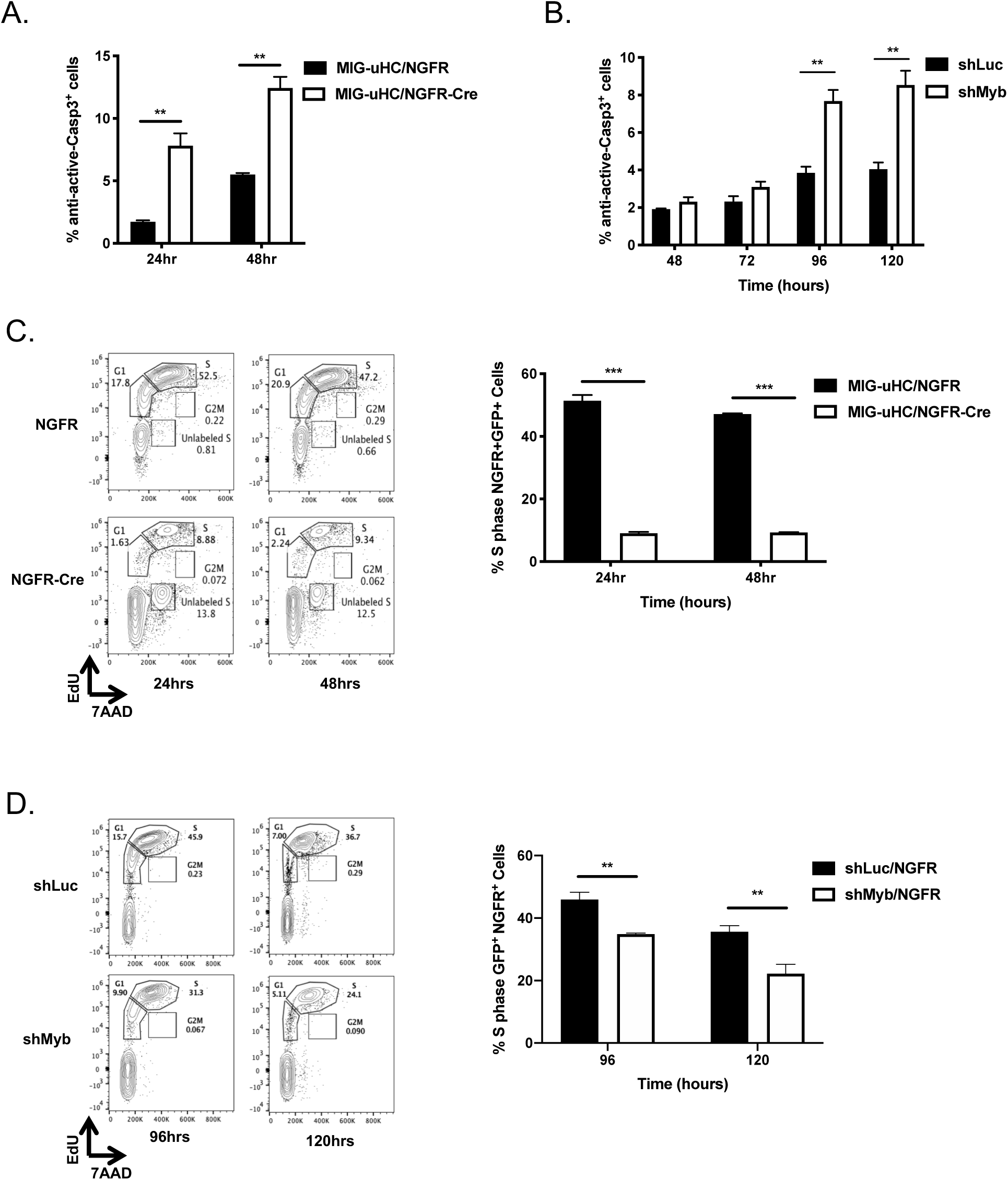
c-Myb Deletion in *Myb*^*ff*^*Rag2*^*-/-*^ *uHC*-Transduced Pro-B cells Leads to Decreased Pre-B Cell Survival and Proliferation. (A and C) Pre-B cells were generated by retrovirus transduction of *Myb*^*ff*^ *Rag2*^*-/-*^ pro-B cells as described in Fig. 1. Following retrovirus transduction, pre-B cells were cultured with 10ng/ml IL-7 and every 24 h, co-transduced GFP^+^ NGFR^+^ cells were analyzed for cell survival and cell cycle distribution. (B and D) *Irf4/8*^*-/-*^ large pre-B cells were transduced with shLuc-GFP or shMyb-GFP and cultured with 10ng/ml IL-7. At the indicated time points, GFP^+^ cells were analyzed for cell survival and cell cycle distribution. (**A**) The proportion of co-transduced GFP^+^ NGFR^+^ active Caspase 3^+^ pre-B cells was analyzed 24 and 48 h post-transduction by flow cytometry. Data are representative of 2 independent experiments. (**B**) The proportion of GFP^+^ active Caspase 3^+^ *Irf4/8*^*-/-*^ large pre-B cells was analyzed 48, 72, 96, and 120 h post-transduction by flow cytometry. Data are representative of 2 independent experiments. (**C**) At 24 h and 48 h post-transduction pre-B cells were labeled with EdU for 2 h and cell cycle analysis was performed by flow cytometry after staining with the Click-iT Plus EdU reaction cocktail to assess DNA synthesis and 7AAD to assess DNA content. The proportion of GFP^+^ NGFR^+^ S-phase cells was determined at each time point. Representative EdU versus 7AAD plots for each condition and time point are shown. Data are representative of 2 independent experiments. (**D**) At 96 h and 120 h post-transduction *Irf4/8*^*-/-*^ cells were labeled with EdU for 2 h and cell cycle analysis was performed by flow cytometry after staining with the Click-iT Plus EdU reaction cocktail and 7AAD. The proportion of GFP^+^ S-phase cells was determined at each time point. Data are representative of 3 independent experiments. All retrovirus transductions were performed with three replicates per condition. *p < 0.05, **p < 0.005, ***p < 0.0005.

### c-Myb-dependent gene expression changes in large pre-B cells include an Ikaros-dependent pro-differentiation gene signature

We have previously reported that rapid cell death after acute deletion at the *Myb*^*f/f*^ locus can limit detection of some genes that are regulated by c-Myb (30). To avoid this problem, we compared gene expression in *Irf4/8*^*-/-*^ large pre-B cells 72 h after transduction with shMyb or shLuc as this is the first time point where we detected a statistically significant decrease in relative recovery of shMyb transduced cells (Fig. 1C). As a result, 5,895 genes were significantly changed with a false discovery rate of p<0.05, 2,979 genes had a log2 fold change > 1, and 725 genes met both of these thresholds, demonstrating that loss of c-Myb expression has broad effects on global gene expression (Fig. 4A) which is consistent with other reports (47–49). To identify major processes that were impacted by the loss of c-Myb, we performed gene set analysis utilizing KEGG pathway and biological process gene ontology (GO-BP) gene sets and validated a panel of key genes involved in these processes by qRT-PCR (Figs. 4B, 4C). Consistent with our phenotypic data, we identified pathways and processes related to proliferation that were enriched in downregulated genes. These downregulated genes included a number of cyclins and cyclin-dependent kinases (Cdks) that drive cell cycle progression. In particular, *Cdk4* (Cdk4) and its target *Ccnd3* (cyclin D3), which is required for the proliferation of large pre-B cells (Supplemental Figure. 2A). We also detected decreased mRNA expression of the transcription factor c-Myc, which is important both to promote transcription of cell cycle genes as well as repress transcription of cell cycle inhibitors. In particular, the cell cycle inhibitor and c-Myc target *Cdkn1b* (p27 or Kip1) was upregulated in our analysis (15, 18, 50). Interestingly, we also found that downregulated genes were enriched in metabolic pathways and processes needed to maintain cell growth and division (Supplemental Figures. 2B, 2C). In particular, expression of the glucose transporter *Slc2a1* (Glut1), and the hexokinase isoform *Hk1* (HK1), which mediates the initial step of glucose processing in the cell, were significantly decreased. In addition, expression of the Glut1 inhibitor *Txnip* (thioredoxin interacting protein), was significantly increased upon loss of c-Myb. *Txnip* inhibits transcription of Glut1 mRNA and promotes Glut1 internalization and proteasomal degradation, suggesting that c-Myb controls the initial steps of glucose metabolism in large pre-B cells (51, 52). Consistent with this notion downregulated genes were significantly enriched in metabolic pathways downstream of glucose uptake that utilize products of glucose catabolism. These metabolic pathways included glycolysis, the biosynthesis of macromolecules such as nucleic acids and fatty acids, and the production of pyruvate which feeds into the TCA cycle and mitochondrial oxidative phosphorylation (8). Thus, analysis of gene sets enriched in downregulated genes suggests that c-Myb might have a previously unappreciated role in the regulation of glucose metabolism. This role would be particularly significant in large pre-B cells, which rely heavily on glucose to fuel their rapid proliferation, and thus is likely to be important to the phenotype of c-Myb-deficient large pre-B cells.

**FIGURE 4.**
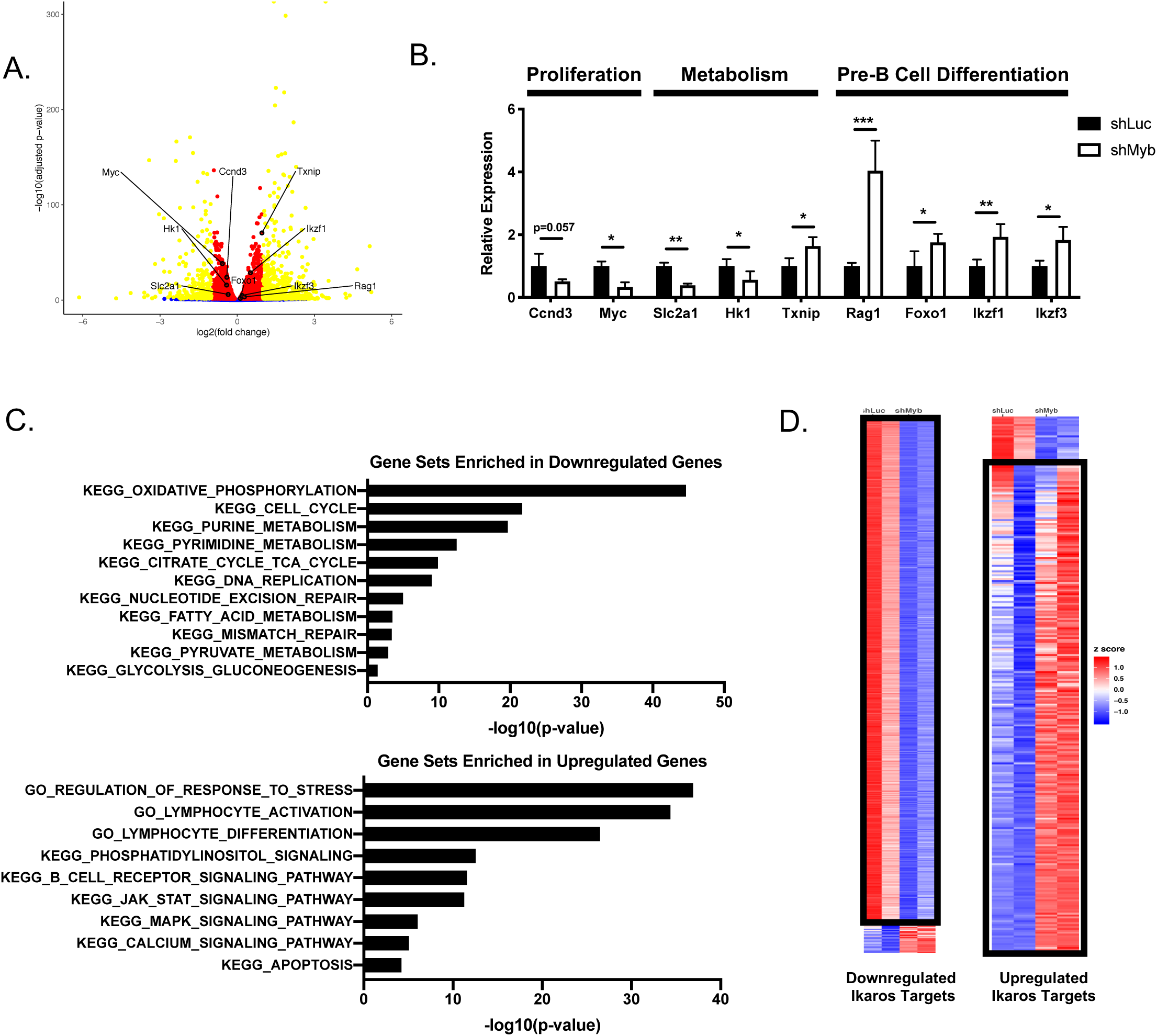
c-Myb-Dependent Gene Expression Changes in Large Pre-B Cells Correlate with an Ikaros-Dependent Gene Signature. *Irf4/8*^*-/-*^ large pre-B cells were transduced with shLuc-GFP or shMyb-GFP and cultured with 10ng/ml IL-7. At 72 h post-transduction, GFP^+^ cells were electronically sorted and total RNA was prepared for genome-wide gene expression profiling by RNA-seq. Retroviral transduction and RNA-seq was performed on 2 replicates per condition. (**A**) Volcano plot of differentially expressed (DE) genes displayed as fold change (log2FC; x-axis) versus statistical significance (p-value; y-axis). Each gene is represented as one dot and is colored based on a threshold adjusted p-value (padj) < 0.05 (red), log2FC > 1 (blue), and the genes that satisfy both of these parameters (yellow). (**B**) Gene set analysis of DE genes with a padj < 0.0001 utilizing KEGG pathway and GO-BP ontology gene sets. The analysis was divided between downregulated and upregulated DE genes. (**C**) qRT-PCR validation of selected DE genes from GFP^+^ *Irf4/8*^*-/-*^ cells electronically sorted at 72 h post-transduction. Gene expression was normalized to expression of *HPRT* and genes were analyzed in triplicate per condition. *p < 0.05, **p < 0.005, ***p < 0.0005. (**D**) Heatmap depicting merged gene signatures of DE genes from shLuc/shMyb transduced *Irf4/8*^*-/-*^ large pre-B cells and direct Ikaros targets differentially expressed at the large to small pre-B cell transition as reported by Ferreiros-Vidal et al. (2013). The overlap in genes that are downregulated or upregulated in both the c-Myb-dependent and Ikaros-dependent gene signatures are indicated by black boxes.

Pathways and processes related to cell stress and apoptosis and, interestingly to B cell activation and/or differentiation including BCR signaling, MAPK signaling, PI3K signaling, and calcium flux were enriched in upregulated genes (Supplemental Figure. 2D). Notably, key genes associated with differentiation from large to small pre-B cells were upregulated including *Foxo1* and *Rag1* (Fig.4B), which are important for the initiation of VJ recombination. In addition, the transcription factor *Irf4* and its downstream targets *Ikzf1* and *Ikzf3* (Fig. 4B), which mediate many of the gene expression changes needed for the large to small pre-B cell transition. Furthermore, we detected increased expression of *Sh2b3* which is a direct Ikaros target gene that inhibits IL-7R signaling through interaction with the proximal kinase Jak3 (2, 22, 53, 54). Overall, the results of gene set analysis suggest that shMyb-transduced large pre-B cells are receiving opposing signals from the downregulation of genes involved in cell growth, proliferation, and metabolism, which typically accompany differentiation to small pre-B cells, and the upregulation of genes promoting cell activation, which would typically increase these same cell growth and proliferation pathways. As we also observe upregulation of cell stress and apoptosis associated genes, these results suggest that a failure to reconcile conflicting gene expression changes might increase cell stress and, if unresolved, could ultimately result in cell death.

Our RNA-seq differential gene expression data and gene set analysis suggested that c-Myb expression is important for maintaining the expression of genes involved in proliferation and metabolism while suppressing the expression of genes involved in transition to the small pre-B cell stage such as the transcription factors Irf4, Ikaros, and Aiolos that were upregulated upon *Myb* knockdown. Downstream of pre-BCR signaling, Irf4 induces expression of Ikaros and Aiolos that mediate gene expression changes important for the large to small pre-B cell transition. In particular Ikaros and Aiolos-mediated gene expression changes lead to decreased metabolism, exit from the cell cycle, and initiation of changes in chromatin structure necessary for VJ recombination at the IgL locus (21). Thus, c-Myb-mediated repression of Ikaros and Aiolos expression could be responsible for many of the gene expression changes related to proliferation, metabolism, and differentiation we detect upon c-Myb knockdown in large pre-B cells. In fact, comparison of c-Myb-dependent differentially expressed genes with a previously published Ikaros footprint revealed that approximately 70% of the direct Ikaros targets that are important for the large to small pre-B cell transition were also changed by c-Myb knockdown (Fig. 4D) (22). Included among these genes were those we had already identified above as genes that have crucial roles in the proliferation (*Ccnd3* and *Myc*), metabolism (*Slc2a1* and *Txnip*), and differentiation (*Foxo1* and *Sh2b3*) of large pre-B cells. We noted that *Hk1*, a critical enzyme mediating the initial step in glucose utilization, was not included in the Ikaros footprint. This result suggested the failure to repress Ikaros and Aiolos expression in c-Myb-deficient large pre-B cells could lead to decreased metabolism, decreased proliferation, and trigger the implementation of a gene expression program promoting differentiation into small pre-B cells.

### c-Myb is critical for large pre-B cell glucose uptake and hexokinase activity

The increased biosynthetic demands of large pre-B cells are predominantly met by increased glucose uptake and processing through glycolysis, which provides critical intermediates for the production of macromolecules needed for cell division (8). Our RNA-seq gene set analysis suggested that c-Myb knockdown in large pre-B cells would result in substantial defects in glucose metabolism. In particular, mRNA expression of Glut1 (*Slc2a1*) and hexokinase 1 (*Hk1*) were significantly decreased upon loss of c-Myb (Fig. 4B). Failure to take up and phosphorylate sufficient glucose due to loss of Glut1 and Hk1 expression would preclude glucose utilization in downstream metabolic pathways. To determine whether c-Myb is important for glucose uptake in large pre-B cells, we transduced *Irf4/8*^*-/-*^ large pre-B cells with shLuc or shMyb and 72 h later incubated them with the fluorescent glucose analog, 2-deoxy-2-[(7-nitro-2,1,3-benzoxadiazol-4-yl) amino]-D-glucose (2-NBDG) (Fig. 5A). By 72 h post-transduction, the time point at which we begin to detect a significant decrease in relative recovery of shMyb transduced *Irf4/8*^*-/-*^ large pre-B cells, we detected an approximately 50% decrease in 2-NBDG uptake, suggesting that the c-Myb-dependent decrease in Glut1 mRNA expression resulted in decreased glucose transport into the cell. To determine if c-Myb is also important for glucose processing, we measured hexokinase activity of shMyb-transduced *Irf4/8*^*-/-*^ large pre-B cells and found, similar to glucose uptake, that c-Myb knockdown resulted in a 50% decrease in overall hexokinase activity by 72 h post-transduction (Fig. 5B). Thus, c-Myb is important for maintaining expression of critical upstream components of glucose metabolism involved in glucose uptake (Glut1) and utilization (Hk1) in large pre-B cells.

**FIGURE 5.**
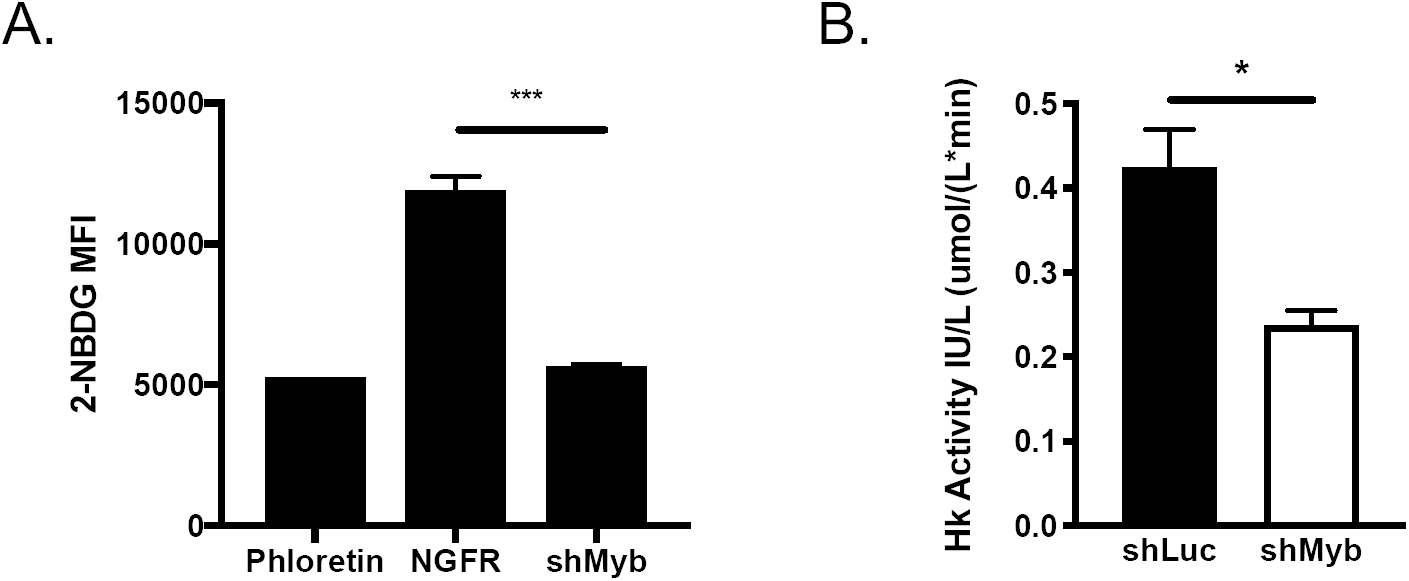
c-Myb Knockdown Leads to Defects in Large Pre-B Cell Glucose Uptake and Hexokinase Activity. *Irf4/8*^*-/-*^ large pre-B cells were transduced with NGFR or NGFR-shMyb (A) or shLuc-GFP or shMyb-GFP (B) and cultured for 72 h with 10ng/ml IL-7. (**A**) At 72 h post-transduction, NGFR^+^ cells were incubated 30 min with the 2-NBDG glucose uptake mix and 2-NBDG MFI was measured by flow cytometry. The glucose uptake inhibitor phloretin was used as a negative control for 2-NBDG uptake. Data is representative of 2 independent experiments. (**B**) At 72 h post-transduction GFP^+^ cells were electronically sorted and protein lysate was prepared for analysis of hexokinase enzymatic activity. Enzyme activity was determined by colorimetric assay and absorbance at 492nm. Data is representative of 2 independent experiments. All retrovirus transductions were performed with three replicates per condition. N = 3, *p < 0.05, **p < 0.005, ***p < 0.0005.

### Expression of Glut1 or Hk1 rescues relative recovery, survival, and proliferation of c-Myb-deficient large pre-B cells

To determine the contribution of c-Myb mediated regulation of glucose uptake/utilization to the c-Myb dependent large pre-B cell phenotype, *Irf4/8*^*-/-*^ large pre-B cells were co-transduced with GFP-shLuc or GFP-shMyb and either tNGFR (control), tNGFR-Glut1 or tNGFR-Hk1 and the relative recovery of shMyb^+^ NGFR^+^ cells was determined over a 96 h time course (Fig. 6A).

**FIGURE 6.**
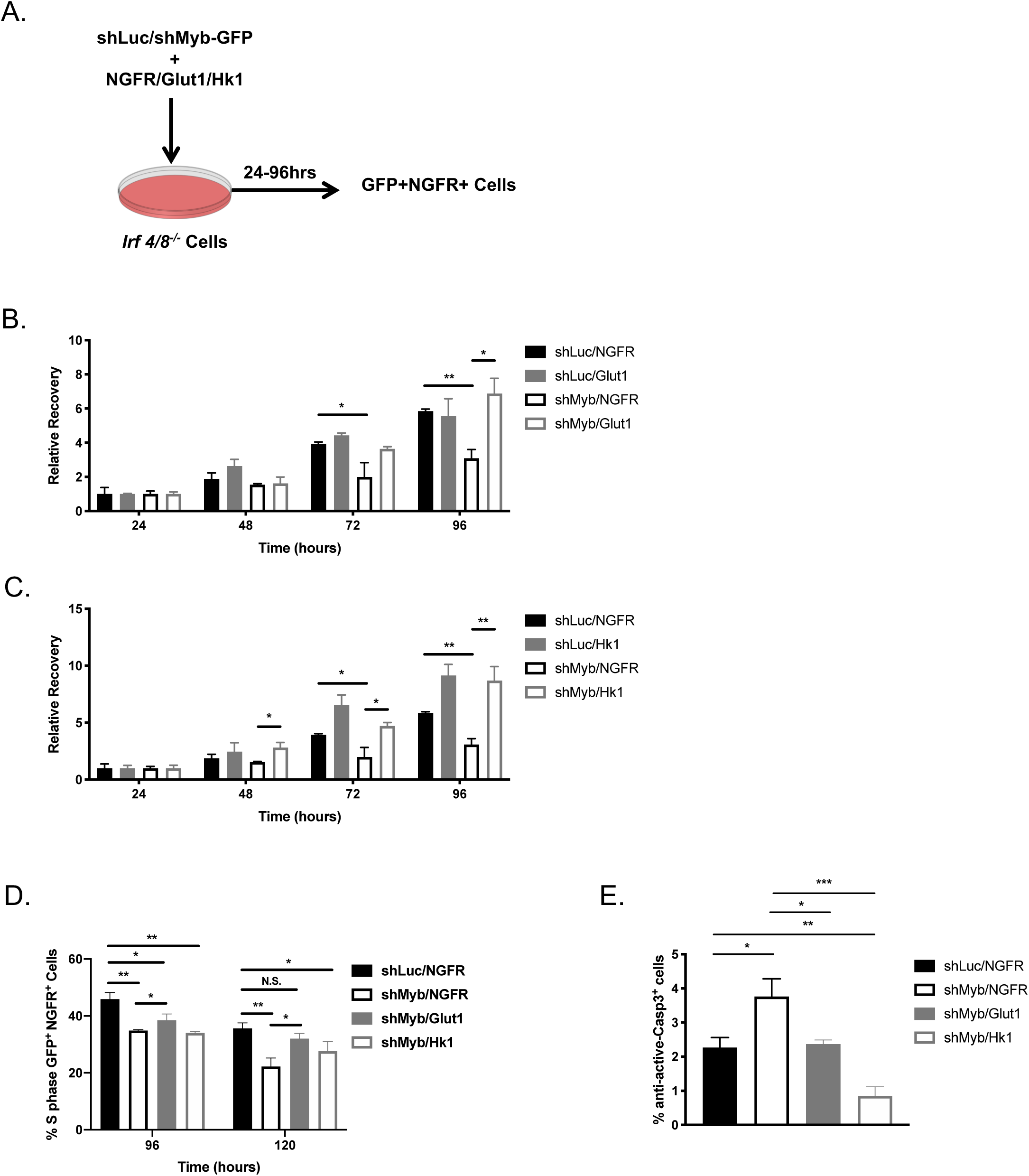
Expression of Glut1 or Hk1 Rescues Recovery and Glucose-Dependent Survival of c-Myb-Deficient Large Pre-B Cells. (**A**) Experimental design used to determine the role of Glut1 and Hk1 expression in large pre-B cells. *Irf4/8*^*-/-*^ large pre-B cells were co-transduced with shLuc-GFP or shMyb-GFP and NGFR, NGFR-Glut1, or NGFR-Hk1 and cultured with 10ng/ml IL-7. Every 24 h post-transduction over a 96 h time course, co-transduced GFP^+^ NGFR^+^ cells were analyzed for total cell numbers and at 96 h post-transduction they were analyzed for cell survival. (B&C) Co-transduced GFP^+^ NGFR^+^ cells were analyzed at 24, 48, 72, and 96 h post-transduction by flow cytometry and relative recovery was determined as described in Figure 1. The numbers of shLuc/shMyb-GFP^+^ cells co-transduced with the NGFR empty vector control were compared to co-transduction with NGFR-Glut1 (**B**) or NGFR-Hk1 (**C**). Data is representative of 2 independent experiments. (**D**) At 96 h and 120 h post-transduction *Irf4/8*^*-/-*^ cells were labeled with EdU for 2 h and cell cycle analysis was performed as described for Fig. 3D. The proportion of GFP^+^ NGFR^+^ S-phase cells was determined at each time point. (**E**) The proportion of co-transduced GFP^+^ NGFR^+^ active Caspase 3^+^ cells was analyzed 96h post-transduction by flow cytometry.

Expression of exogenous Glut1 had little effect on the relative recovery of c-Myb sufficient cells (shLuc/Glut1) (Fig. 6B). However, Glut1 rescued the relative recovery of *Irf4/8*^*-/-*^ large pre-B cells 72 h post-transduction such that the recovery of shMyb/Glut1 transduced cells was comparable to control, shLuc/NGFR transduced cells. Thus, increasing the capacity of *Irf4/8*^*-/-*^ large pre-B cells to take up glucose after c-Myb knockdown is sufficient to restore their relative recovery.

Transduction with Hk1 increased the relative recovery of c-Myb-sufficient (shLuc/tNGFR) *Irf4/8*^*-/-*^ large pre-B cells by approximately 1.5-fold suggesting that Hk1 overexpression increased the efficiency of glucose utilization resulting in increased relative recovery of *Irf4/8*^*-/-*^ large pre-B cells (Fig. 6C). This is consistent with the high affinity of Hk1 for glucose and greater activity as compared to other hexokinase isoforms (55). Importantly, expression of exogenous Hk1 resulted in increased recovery of *Irf4/8*^*-/-*^ large pre-B cells after c-Myb knockdown (shMyb/HK1) to a level comparable to that of shLuc/tNGFR or shLuc/Hk1 transduced control cells. Thus, forced expression of the key mediators of glucose uptake and utilization, Glut1 and Hk1, completely rescued the relative recovery resulting from c-Myb knockdown in *Irf4/8*^*-/-*^ large pre-B cells. In addition, the increased relative recovery induced by exogenous Hk1 expression as compared to Glut1 expression suggests that Hk1 activity is the limiting factor in the ability of large pre-B cells to utilize glucose. Overall, this result indicates that c-Myb-dependent regulation of glucose uptake and utilization through hexokinase activity are crucial to maintain the large pre-B cell compartment.

To better characterize the Glut1 and Hk1-mediated rescue of c-Myb-deficient *Irf4/8*^*-/-*^ large pre-B cells, we assessed proliferation by EdU uptake and 7AAD DNA staining at 96 h and 120 h after co-transduction (Fig. 6D). Expression of Glut1 in c-Myb-deficient *Irf4/8*^*-/-*^ large pre-B cells (shMyb/Glut1) significantly increased the proportion of S-phase cells over shMyb-transduced controls at both 96 h and 120 h post-transduction. In addition, at 120 h post-transduction, the proportion of S-phase shMyb/Glut1 transduced cells was restored to the level of shLuc/NGFR transduced control cells. Thus, exogenous expression of Glut1 was able to rescue the proliferation defect in c-Myb-deficient *Irf4/8*^*-/-*^ large pre-B cells. In contrast, expression of Hk1 in shMyb-transduced *Irf4/8*^*-/-*^ large pre-B cells (shMyb/Hk1) failed to significantly increase the proportion of S-phase cells over shMyb-transduced controls at either 96 h or 120 h post-transduction, although there appeared to be a slight increase in S-phase cells at the 120 h time point. Thus, in the context of c-Myb knockdown, exogenous expression of Glut1 but not Hk1 was able to restore *Irf4/8*^*-/-*^ large pre-B cell proliferation. This result suggested that expression of Hk1 rescued relative recovery of c-Myb-deficient *Irf4/8*^*-/-*^ large pre-B cells through a mechanism other than promoting proliferative expansion.

In addition to fueling metabolism and proliferation, glucose uptake and utilization through hexokinase activity is critical for cell survival (52, 56). Hk1 associates with mitochondria and interferes with binding of pro-apoptotic Bcl-2 family members, maintains mitochondrial membrane integrity, and prevents the release of cytochrome c associated with apoptosis in the FL5.12 lymphoid cell line. To determine if exogenously supplied Glut1 or Hk1 could decrease apoptosis in *Irf4/8*^*-/-*^ large pre-B cells after c-Myb knockdown, the proportion of active Caspase 3^+^ cells was assessed in *Irf4/8*^*-/-*^ large pre-B cells 96 h after co-transduction (Fig. 6E). While *Irf4/8*^*-/-*^ large pre-B cells transduced with Glut1 decreased the proportion of active Caspase 3^+^ cells after c-Myb knockdown to a proportion comparable to control cells (GFP/tNGFR), expression of Hk1 resulted in an even further decrease in active Caspase 3^+^ cells. Taken together, these results suggest that in the context of c-Myb-deficiency, increasing the availability of glucose through expression of Glut1 is able to restore both proliferation and survival of *Irf4/8*^*-/-*^ large pre-B cells. In contrast, the function of exogenous Hk1 expression in c-Myb-deficient *Irf4/8*^*-/-*^ large pre-B cells, which exhibit decreased expression of Glut1 and decreased glucose uptake, may be focused toward preventing apoptosis over promoting proliferative expansion. Thus, these results indicate that the increase in relative recovery we observe upon expression of Hk1 is largely due to increased cell survival. Furthermore, our observation that expression of either Glut1 and Hk1 rescues apoptotic cell death suggests that decreased cell survival constitutes the dominant phenotype of c-Myb-deficient *Irf4/8*^*-/-*^ large pre-B cells.

### Ikaros-dependent and Ikaros-independent functions of c-Myb in large pre-B cells

To transition from cycling large pre-B cells to quiescent small pre-B cells, large pre-B cells exit the cell cycle and significantly decrease metabolic activity (2, 3). The transcription factor Ikaros and its family member Aiolos mediate most of the gene expression changes needed for the large to small pre-B cell transition and we have found, approximately 70% of the Ikaros targets important for this transition were also differentially expressed upon c-Myb knockdown in *Irf4/8*^*-/-*^ large pre-B cells (Figure 4D). Many of the gene expression changes induced by both c-Myb knockdown and Ikaros are also critical for mediating the phenotype we have observed in c-Myb-deficient large pre-B cells including decreased proliferation as well as decreased glucose uptake/utilization. Furthermore, we have demonstrated that exogenous expression of genes important for glucose uptake/utilization, Glut1 and Hk1, rescues survival of shMyb-transduced *Irf4/8*^*-/-*^ large pre-B cells (Figure 6D). These results suggest that failures to repress Ikaros expression in c-Myb-deficient large pre-B cells could account for the defects in proliferation, metabolism, and survival we have observed upon c-Myb knockdown. Therefore, we determined whether inhibition of Ikaros activity could rescue defects in recovery of large pre-B cells after c-Myb knockdown.

Ikaros undergoes alternative splicing to produce isoforms with variable DNA binding capability. Isoforms of Ikaros that are unable to bind DNA have a dominant negative effect on Ikaros function (57, 58). This occurs by dimerization with full length isoforms of Ikaros family members to sequester them and prevent DNA binding as well as through competition with full-length Ikaros isoforms for participation in chromatin remodeling complexes. To determine if repression of Ikaros and Aiolos could rescue the decreased relative recovery that we observed upon c-Myb knockdown in *Irf4/8*^*-/-*^ large pre-B cells, we utilized a DNA binding domain point-mutant of Ikaros (Ik159A), which acts similarly to the naturally occurring dominant negative Ikaros isoforms (22). *Irf4/8*^*-/-*^ large pre-B cells were co-transduced with tNGFR/MIG-R1, tNGFR/MIG-IK159A, shMyb/MIG-R1 or shMyb/MIG-Ik159A and the relative recovery of each co-transduced pair was assessed every 24 h over a 96 h time course (Fig. 7A). Expression of the Ik159A mutant had little or no effect on the relative recovery of c-Myb-sufficient cells (compare tNGFR/MIG-R1 with tNGFR/MIG-Ik159A) while shMyb/MIG-R1 resulted in approximately a 60% and 70% decrease in relative recovery at 72 h and 96 h post-transduction respectively (Fig. 7B). However, while we did detect a small increase in relative recovery of shMyb/MIG-Ik159A transduced cells, this was not sufficient to fully restore relative recovery to a level comparable to tNGFR/MIG-R1 transduced controls. Thus, inhibition of Ikaros activity using the Ik159A mutant is unable to account for the entire phenotype induced by c-Myb knockdown in *Irf4/8*^*-/-*^ large pre-B cells, suggesting that there are Ikaros dependent and independent changes in gene expression that contribute to the decreased proliferative expansion in this model.

**FIGURE 7.**
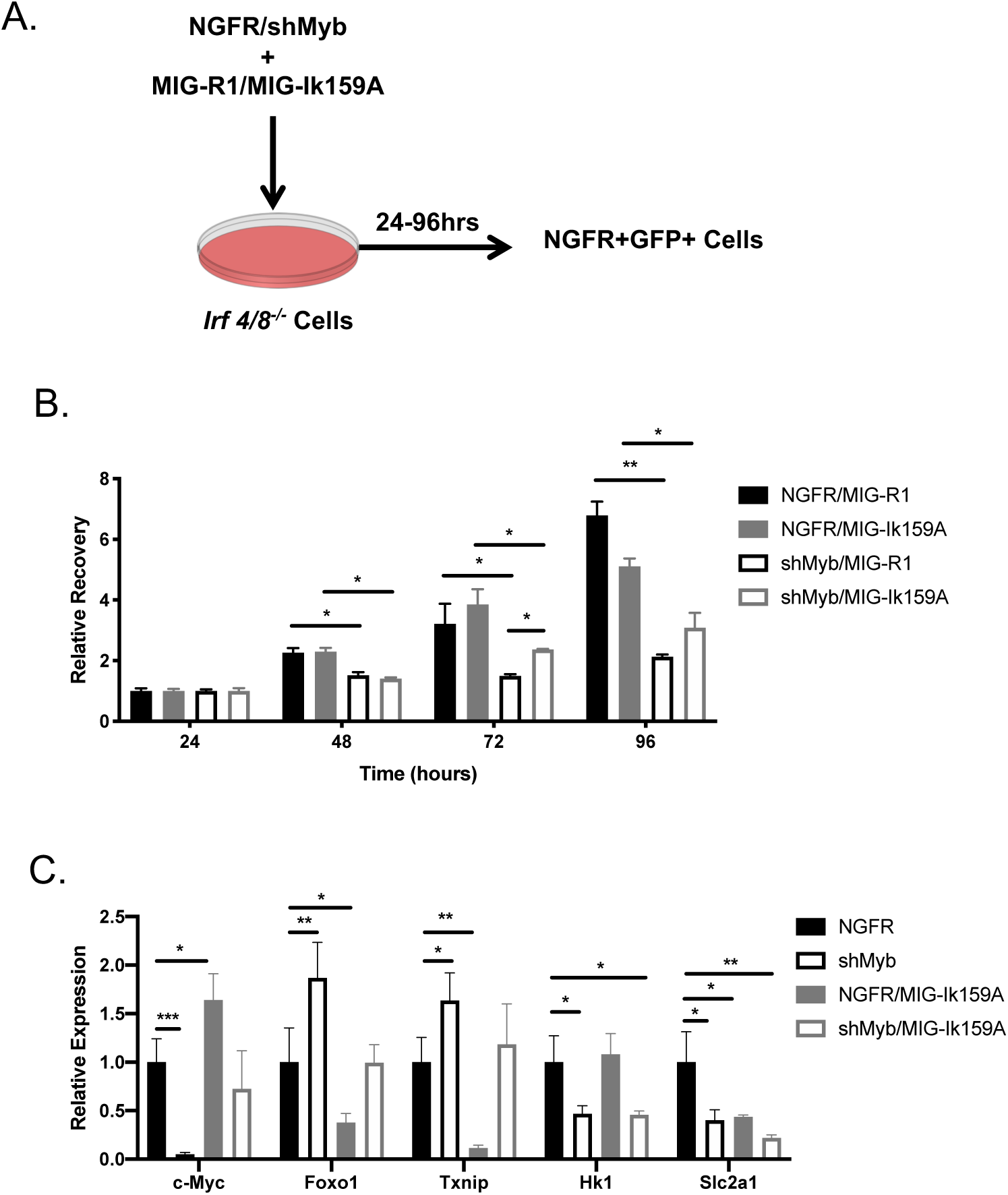
Ikaros-Dependent and –Independent gene expression mediated by c-Myb in Large Pre-B Cells. (**A**) Experimental design used to determine the role of dominant negative mutant Ikaros (Ik159A) expression in large pre-B cells. *Irf4/8*^*-/-*^ large pre-B cells were co-transduced with NGFR or NGFR-shMyb and MIG-R1 or MIG-Ik159A and cultured with 10ng/ml IL-7. Every 24 h post-transduction over a 96 h time course, co-transduced NGFR^+^ GFP^+^ cells were analyzed for total cell numbers and at 72 h post-transduction gene expression changes were analyze. (**B**) Co-transduced NGFR^+^ GFP^+^ cells were analyzed at 24, 48, 72, and 96 h post-transduction by flow cytometry and relative recovery was determined as described in Fig. 1. The numbers of NGFR/NGFR-shMyb^+^ cells co-transduced with the MIG-R1 empty vector control were compared to co-transduction with MIG-Ik159A. Data is representative of 2 independent experiments. (**C**) At 72 h post-transduction single-transduced NGFR^+^ and co-transduced NGFR^+^ GFP^+^ cells were electronically sorted and total RNA was prepared for qRT-PCR. Gene expression was normalized to expression of *HPRT* and genes were analyzed in triplicate per condition. *p < 0.05, **p < 0.005, ***p < 0.0005.

Failure of the dominant negative Ik159A mutant to fully rescue the relative recovery of shMyb transduced *Irf4/8*^*-/-*^ large pre-B cells suggested that inhibition of Ikaros activity is insufficient to restore all aspects of the c-Myb-dependent large pre-B cell phenotype. Following this result, we considered 2 major possibilities: 1) Ik159A was not expressed in sufficient amounts to sequester Ikaros or 2) an Ikaros-independent role for c-Myb in large pre-B cells prevented complete recovery after Ikaros neutralization. To address these possibilities, we determined if Ik159A could restore Ikaros-dependent gene expression changes in large pre-B cells following c-Myb knockdown. *Irf4/8*^*-/-*^ large pre-B cells were co-transduced with tNGFR, tNGFR-shMyb, tNGFR/MIG-Ik159A or shMyb/MIG-Ik159A and both tNGFR^+^ and tNGFR^+^ GFP^+^ cells were electronically sorted at 72 h post-transduction. We then determined mRNA expression of a panel of genes differentially expressed upon c-Myb knockdown in *Irf4/8*^*-/-*^ large pre-B cells that have been reported to be Ikaros targets. In each case we would expect the Ik159A mutant to reverse expression mediated by Ikaros upon c-Myb knockdown (Fig. 7C) (22).

c-Myc mRNA expression was downregulated upon c-Myb knockdown in *Irf4/8*^*-/-*^ large pre-B cells (Fig. 7C) and is repressed by direct Ikaros binding (50). We found that transduction with the Ik159A mutant resulted in increased *Myc* mRNA expression in c-Myb-sufficient *Irf4/8*^*-/-*^ large pre-B cells, consistent with a dominant negative function in counteracting Ikaros-mediated repression of *Myc*. Furthermore, co-transduction with shMyb and the Ik159A mutant resulted in a level of *Myc* mRNA expression comparable to the control (c-Myb-sufficient *Irf4/8*^*-/-*^ cells), indicating that Ik159A is able to inhibit Ikaros function to oppose the decreased expression of *Myc* resulting from c-Myb knockdown and the resultant increase in Ikaros. *Foxo1* and *Txnip* mRNA expression are positively regulated by direct Ikaros binding (22) and were also upregulated upon c-Myb knockdown in *Irf4/8*^*-/-*^ large pre-B cells (Fig. 7C). Transduction with the Ik159A mutant produced the opposite result and resulted in increased *Foxo1* and *Txnip* mRNA expression in c-Myb sufficient *Irf4/8*^*-/-*^ large pre-B cells. Again, co-transduction with shMyb and the Ik159A mutant resulted in *Foxo1* and *Txnip* mRNA expression comparable to control *Irf4/8*^*-/-*^ cells. Thus, *Myc, Foxo1* and *Txnip* mRNA appear to be controlled in *Irf4/8*^*-/-*^ large pre-B cells by Ikaros in a c-Myb dependent fashion, demonstrating that the dominant negative Ik159A was able to reverse c-Myb-dependent gene expression changes in *Irf4/8*^*-/-*^ large pre-B cells. As expected, *Slc2a1* (Glut1) expression, which is repressed by direct Ikaros binding (22, 59), was also repressed by transduction with shMyb (Fig. 7C). However, unexpectedly, expression of *Slc2a1* was also repressed after transduction with Ik159A. Thus, it appears that Ikaros mediates repression of Glut1 in a manner that is unaffected by interaction with the dominant negative Ik159A mutant. Taken together, these results are in agreement with a previous study (22) that examined Ikaros dependent gene expression in the mouse B3 pre-B cell line (Supplemental Figure 3).

In contrast to the genes regulated by c-Myb and Ikaros, Hk1 mRNA expression was decreased with c-Myb knockdown, but was unaffected by Ik159A alone or when co-transduced with shMyb, suggesting that *Hk1* is largely regulated by c-Myb in an Ikaros-independent manner. This is consistent with previous reports that Ikaros has no or little effect on Hk1 expression (22, 59) as summarized in Supplemental Figure 3. Thus, c-Myb has both Ikaros dependent and independent effects on genes in large pre-B cells. Taken together, our results demonstrate that c-Myb plays a crucial role in balancing the expression of genes that are important for maintaining the large pre-B cell compartment while suppressing the premature expression of genes that are important for quiescence and differentiation to the small pre-B cell compartment.

## Discussion

c-Myb is crucial for pro-B cell survival as well as transition to the large pre-B cell compartment (28, 29). However, it was not known if c-Myb plays additional roles within the pre-B cell compartment. Rapidly cycling large pre-B cells are dependent on glucose uptake and the production of glycolytic intermediates to facilitate cell division (4, 5). We demonstrate that c-Myb plays a crucial and unappreciated role in regulating glucose uptake and utilization in the large pre-B cell compartment that is critical to maintain proliferation and prevent apoptosis. c-Myb promotes glucose metabolism in large pre-B cells at the level of both glucose uptake and utilization through hexokinase activity. c-Myb regulates glucose uptake through the glucose transporter Glut1. Loss of c-Myb leads to decreased expression of Glut1 mRNA as well as increased expression of the Glut1 inhibitor Txnip. Txnip has multiple metabolism-related functions including inhibiting Glut1 by promoting its internalization from the plasma membrane to the lysosome and subsequent proteasomal degradation as well as regulating Glut1 mRNA levels (60–62). Downstream of glucose uptake, c-Myb regulates glucose phosphorylation through the glycolytic enzyme Hk1, which mediates the initial step of glucose metabolism, conversion of glucose into glucose-6-phosphate for downstream applications such as glycolysis (55). c-Myb knockdown in large pre-B cells leads to decreased Hk1 mRNA expression and a significant reduction in overall hexokinase activity. Notably, c-Myb regulation of the initial steps in glucose metabolism is critical for the c-Myb-dependent large pre-B cell phenotype as exogenous expression of Glut1 or Hk1 is able to fully restore recovery of large pre-B cells after c-Myb knockdown. We further demonstrate that c-Myb represses mRNA expression of Ikaros, which is critical to prevent the premature initiation of a gene expression program that promotes differentiation to the small pre-B cell compartment.

Taken together, our results demonstrate an important role for c-Myb in maintaining homeostasis across the pre-BCR checkpoint by controlling key mediators of glucose uptake and utilization while preventing premature differentiation by repressing Ikaros expression.

### Role of glucose metabolism in large pre-B cell survival

Control of glucose metabolism by c-Myb in large pre-B cells has critical implications for cell survival. Proper glucose metabolism is crucial to maintain mitochondrial integrity and ultimately to prevent apoptotic cell death. In particular, growth factor receptor signaling through Akt has been reported to promote survival of the FL5.12 pro-B cell line and Rat1a fibroblasts through a mechanism dependent on glucose uptake and hexokinase activity (52, 56). The hexokinase isoforms Hk1 and Hk2, also known as mitochondrial hexokinases (mtHk), are able to associate with the outer mitochondrial membrane (63, 64). Furthermore, mtHks largely utilize intramitochondrial sources of ATP to catalyze glucose phosphorylation and thus couple glycolysis and oxidative phosphorylation. Akt signaling has been reported to inhibit dissociation of mtHks from the mitochondrial membrane in Rat1a fibroblasts and mtHk dissociation is correlated with an impaired ability of growth factors and Akt to maintain mitochondrial integrity and inhibit apoptosis (65, 66). This survival pathway appears to be independent of Bcl-2 family member expression as mtHk dissociation induces cytochrome c release even in the absence of the pro-apoptotic proteins Bax and Bak (66). Thus, activation of Akt downstream of IL-7R and pre-BCR signaling coordinates large pre-B cell proliferation and survival in a manner dependent on glucose uptake and hexokinase-mitochondria interaction (56, 66, 67). Our data is consistent with a pro-survival function for mtHKs as we found that in addition to rescuing large pre-B cell relative recovery, exogenous expression of Hk1 also decreased the proportion of active Caspase 3^+^ cells (Fig. 6D), indicating that Hk1 reduces apoptosis and promotes survival of large pre-B cells after loss of c-Myb expression. Therefore, while our RNA-seq and gene set analysis did not indicate that expression of Akt was altered by c-Myb knockdown, defects in glucose uptake and hexokinase activity would compromise the ability of Akt to promote large pre-B cell survival. Thus c-Myb-dependent deficiencies in large pre-B cell glucose metabolism are directly linked to defects in cell survival.

### Ikaros-dependent and Ikaros-independent functions of c-Myb in large pre-B cells

The transcription factor Ikaros and its family member Aiolos are critical for initiating the pro-differentiation gene expression program and associated decreases in proliferation and metabolic activity that occur during the the large to small pre-B cell transition (20, 50, 68). Expression of both Ikaros and Aiolos were increased in *Irf4/8*^*-/-*^ large pre-B cells after c-Myb knockdown and we found that approximately 70% of genes included in a genomic footprint of Ikaros targets that are important for the large to small pre-B cell transition were also differentially expressed upon c-Myb knockdown in large pre-B cells (22). This suggested that failure to repress Ikaros expression could account for the gene expression changes and phenotypic defects we have observed upon c-Myb knockdown in large pre-B cells. We found that expression of a dominant negative Ikaros mutant (Ik159A) was not able to rescue the survival of large pre-B cells after c-Myb knockdown although it was expressed at sufficient amounts to restore expression of direct Ikaros target genes, suggesting genes that are controlled by c-Myb were crucial for the survival of large pre-B cells. We found that Ik159A was not able to restore expression of Hk1 and this is consistent with previous data demonstrating that HK1 is not significantly impacted by Ikaros activity (22). Thus, the phenotype that we have described in large pre-B cells after knockdown of c-Myb is mediated by both Ikaros dependent and independent changes in gene expression.

### Role for c-Myb at the large to small pre-B cell transition

c-Myb expression has long been linked with the immature stages of hematopoietic cell differentiation (24). Early studies utilizing myeloid and erythroid leukemia cell lines associated c-Myb with inhibiting differentiation and maintaining proliferation as c-Myb expression was downregulated by chemically induced differentiation, while forced expression of c-Myb inhibited differentiation (73–76). We have demonstrated that c-Myb has a similar role in large pre-B cells, maintaining them in an immature, proliferating state and inhibiting differentiation, at least in part by repressing expression of Ikaros and Aiolos. At the pre-B cell stage, Ikaros and Aiolos act together to initiate changes in gene expression that are critical for large pre-B cells to exit the cell cycle and differentiate into small pre-B cells (50, 71). Thus, repression of Ikaros and Aiolos expression by c-Myb is consistent with an established role for c-Myb in antagonizing differentiation and promoting proliferation of immature hematopoietic cells. A role for c-Myb in opposing differentiation from large to small pre-B cells would also suggest that c-Myb expression and/or c-Myb activity must be decreased or modified to permit differentiation from large to small pre-B cells. We also noted that c-Myb-deficient large pre-B cells decreased in size but failed to accumulate CD2^+^ small pre-B cells (Fig. 2A), suggesting that c-Myb expression is important to facilitate differentiation into small pre-B cells. However, it is likely the failure to accumulate CD2^+^ pre-B cells represents increased apoptosis due to the Ikaros-independent role for c-Myb in regulating glucose uptake and utilization as well as glucose-dependent survival. In this scenario, despite increased expression of Ikaros and Aiolos, decreased expression of c-Myb fails to permit the accumulation of CD2^+^ pre-B cells and instead, leads to cell death. Thus, it is likely that continued c-Myb expression is required to maintain survival while the shift in pre-BCR signaling that accompanies the large to small pre-B cell transition leads to a sufficient increase in Ikaros expression to overcome c-Myb-mediated repression and thus to permit differentiation. Alternatively, it is possible that increased Irf4 expression, induced by pre-BCR signaling, is sufficient to overcome repression of Ikaros by c-Myb, allowing for expression of genes required for transition to the small pre-B cell compartment yet allowing for sufficient glucose utilization for survival.

### Implications of c-Myb function in normal large pre-B cells for B-ALL

Due to the rapid proliferation of large pre-B cells and the tight regulation needed to coordinate cell cycle exit with differentiation and the initiation of VJ recombination, the pre-B cell stage is particularly prone to the accumulation of transforming mutations that result in the development of B-cell acute lymphoblastic leukemia (B-ALL) (82). Overall, these transforming mutations promote survival and proliferation and inhibit cell cycle arrest and differentiation, perpetually maintaining cells in a large pre-B cell state. We have demonstrated a similar role for c-Myb in normal large pre-B cells, suggesting that c-Myb would have a similar role in leukemias arising at the large pre-B cell stage of development. In fact, c-Myb has been implicated in promoting and maintaining the development of B-ALL in both mouse and human model systems (83, 84). A key hallmark of cancer cells is their altered metabolism, which shifts to aerobic glycolysis and promotes enhanced and sustained growth, proliferation and survival (85). While a role for c-Myb in cancer cell metabolism has not been described, it is interesting to speculate on the implications that a role for c-Myb in maintaining glucose metabolism of normal large pre-B cells would have for understanding c-Myb function in B-ALL. Ikaros has also been implicated in the development of B-ALL and functions as a tumor suppressor that is inactivated in over 80% of cases of BCR-ABL^+^ B-ALL (86, 87). In addition, Ikaros has been demonstrated to exert its tumor suppressor function in B-ALL by acting as a “metabolic gatekeeper” to enforce a chronic state of energy deprivation as a safeguard against autoimmunity and leukemic transformation (88). Therefore, c-Myb regulation of early steps of glucose metabolism as well as c-Myb-mediated repression of Ikaros expression at the large pre-B cell stage have critical implications for B-ALL. Our findings suggest that increased c-Myb could contribute to the development and maintenance of B-ALL through repression of Ikaros expression and prevention of Ikaros-mediated cell cycle exit, metabolic inhibition, and differentiation. Thus, this work provides a rationale for targeting c-Myb or c-Myb-regulated pathways as a therapeutic strategy for cases of B-ALL both as a means of interfering with cancer metabolism as well as to relieve repression of Ikaros and enable its functions as a tumor suppressor and pro-differentiation factor in large pre-B cells.

## Supporting information

Daamen et al Supplemental Figs

## Acknowledgements

We thank the Flow Cytometry Core Facility (FCCF) for cell sorting and the Genome Analysis and Technology Core at the University of Virginia for high throughput sequencing. We would particularly like to thank Joanne Lannigan, Mike Solga and Claude Chew in the FCCF for expert help and advice. The authors are indebted to Dr. Matthias Merkenschlager for a fascinating and helpful discussion. We also thank Drs. Merkenschlager and Hilde Schjerven for providing Ikaros and Ik159A encoding retroviruses and Dr. Jeffrey Rathmell for providing Glut1 and Hk1 encoding retrovirus vectors. We thank Drs. Ulrike M. Lorenz and Loren D. Erickson for comments on the manuscript.

## 3 Abbreviations used in this paper

HSC: hematopoietic stem cell
SOCS: suppressor of cytokine signaling
CISH: cytokine-inducible SH2-containing protein
tNGFR: truncated nerve growth factor receptor
shRNA: short hairpin RNA; CA, constitutively active
qRT-PCR: quantitative real time PCR
ChIP: chromatin immunoprecipitation

## Notes

1 This work was supported in part by National Institutes of Health (NIH) grants, AI059294 and GM100776 (TPB), NIH Training Grant AI07496 (to ARD), Careers in Immunology Fellowship (ARD).

### Competing Interest Statement

The authors have declared no competing interest.

