## Supplementary material for "c-Myb expression is critical to maintain proliferation and glucose metabolism of large pre-B cells": Daamen et al Supplemental Figs

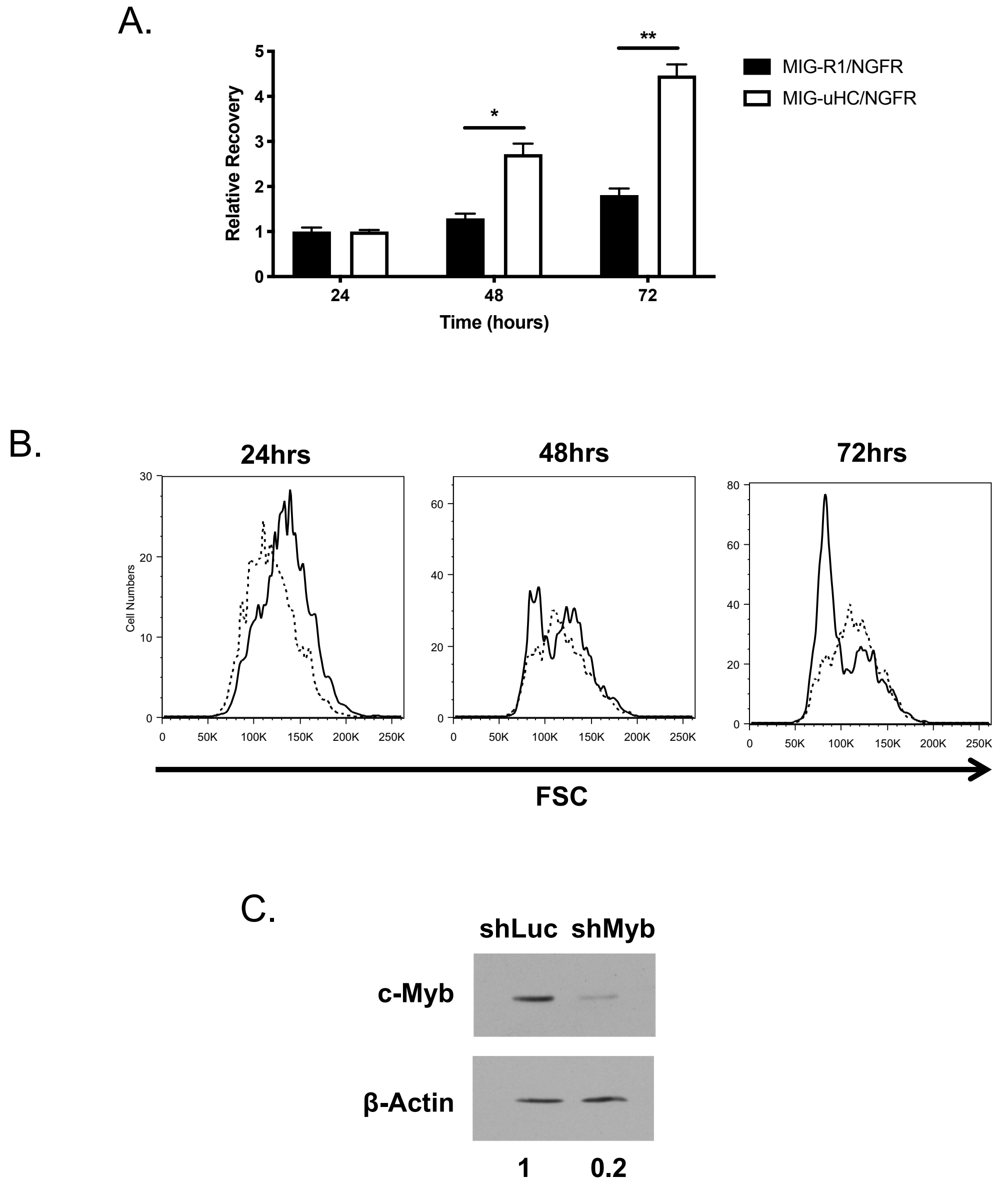


**Supplemental Figure 1. Efficacy of the uHC transduction and *Irf4/8*$^{-/-}$ large pre-B cell models.**
Pro-B cells from *Myb*$^{ff}$ *Rag2*$^{-/-}$ mice were co-transduced with MIG-R1 or MIG-17.2.25 (uHC) and NGFR or NGFR-Cre and cultured with 10ng/ml IL-7. (A) Total numbers of co-transduced GFP$^{+}$ NGFR$^{+}$ cells was analyzed 24, 48, and 72 h post-transduction by flow cytometry. Relative recovery was determined by normalization to the total number of co-transduced cells at the 24 h time point. Retrovirus transductions were done in triplicate. *$p < 0.05$, **$p < 0.005$. (B) Forward scatter (FSC) histograms of co-transduced GFP$^{+}$ NGFR$^{+}$ cells from (A). The solid line represents MIG-uHC/NGFR transduced cells (pre-B) and the dotted line represents MIG-R1/NGFR transduced cells (pro-B). Data is representative of 2 independent experiments. (C) *Irf4/8*$^{-/-}$ large pre-B cells were transduced with shLuc-GFP or shMyb-GFP and cultured with 10ng/ml IL-7. GFP$^{+}$ cells were electronically sorted at 72 h post-transduction and knockdown efficiency of c-Myb protein was analyzed by Western blot. β-Actin was used as a loading control. Data is representative of 3 independent experiments.

#### Supplemental Figure 2

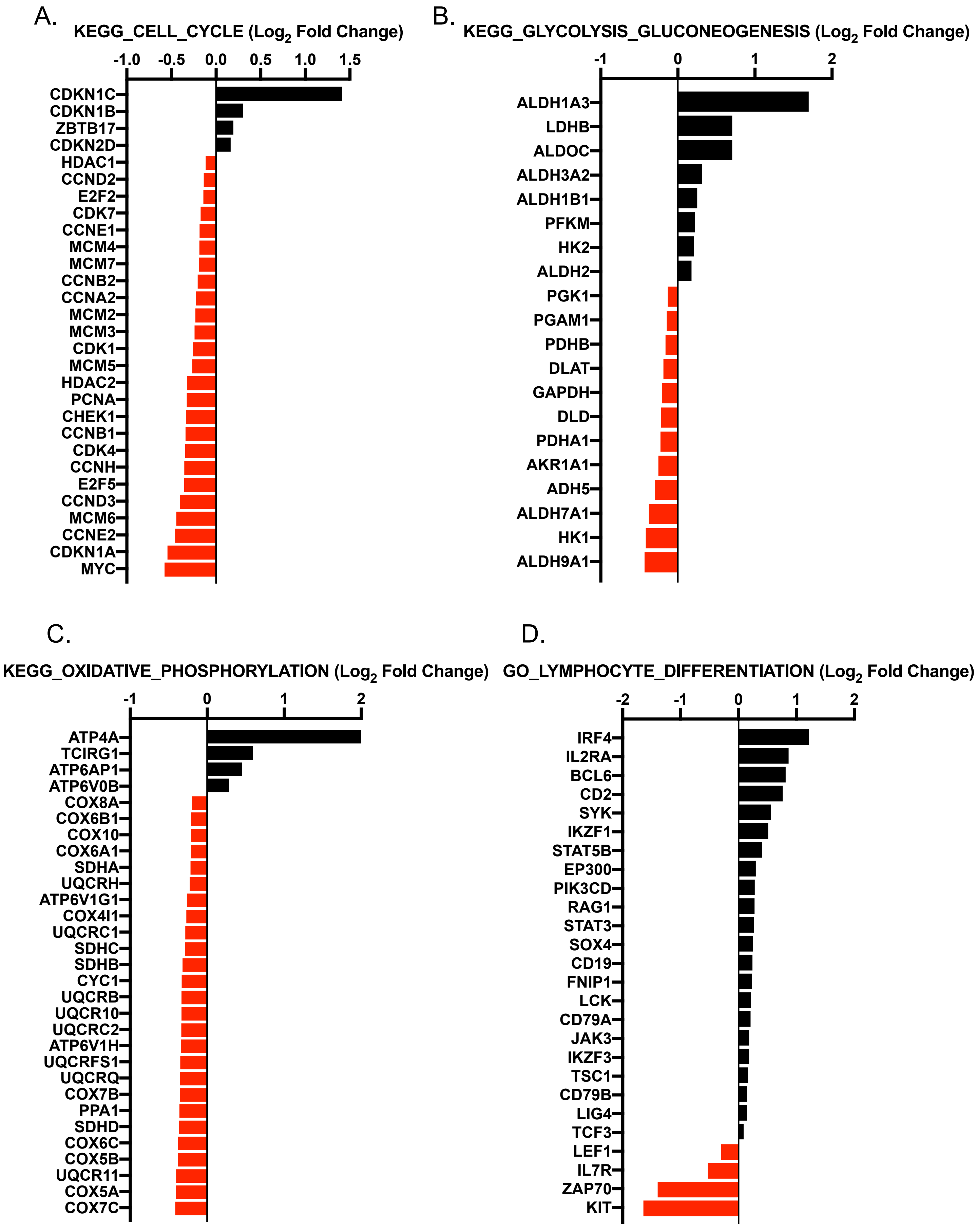

**Supplemental Figure 2. Gene set enrichment of c-Myb-dependent differentially expressed genes.** c-Myb-dependent gene expression changes in large pre-B cells were identified by RNA-seq of *Irf4/8*$^{-/-}$ large pre-B cells after shMyb transduction as described in Fig. 4. Differential expression of genes identified from gene set analysis described in Fig. 4B in the Cell_Cycle (A), Glycolysis_Gluconeogenesis (B), and Oxidative_Phosphorylation (C) KEGG pathway gene sets and the Lymphocyte_Differentiation (D) GO gene set is shown. Genes upregulated upon c-Myb knockdown are shown in black and genes downregulated upon c-Myb knockdown are shown in red.

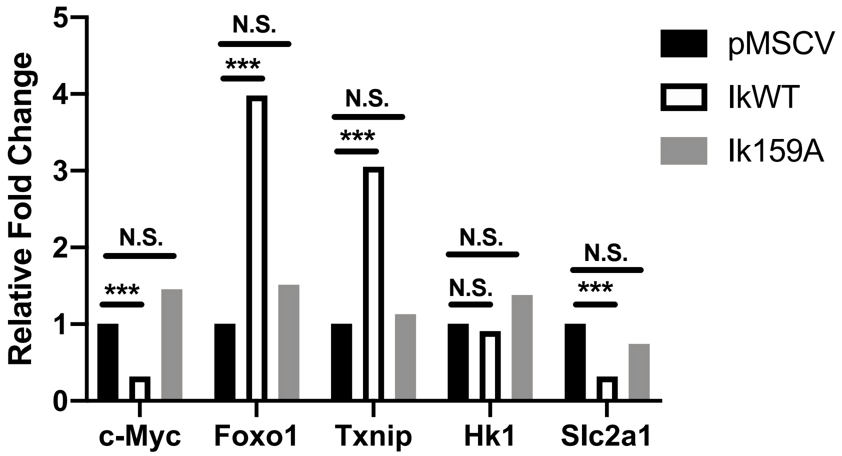


**Supplemental Figure 3. Effects of wild type Ikaros and Ik159A on gene expression in pre-B cells.** Microarray differential expression analysis of B3 pre-B cells transduced with pMSCV, IkWT, or Ik159A was performed as described in Ferreiros-Vidal et al, Blood (2013). Log2 fold change values comparing pMSCV empty vector transduced cells to IkWT or Ik159A transduced cells were converted to fold changes with the empty vector control set as 1. Significance was determined by Benjamini-Hochberg adjusted p-value comparing the pMSCV empty vector to fold changes induced by IkWt or Ik159A transduction.

### Supplemental Table I

| Quantitative Real Time PCR Primers | | |
|---|---|---|
| **Gene** | **Forward Primer** | **Reverse Primer** |
| Ccnd3 | ATGCTGGAGGTGTGTGAGGA | CCACAGCCTGGTCCGTATAG |
| Foxo1 | AAGGATAAGGGCGACAGCAA | TGGATTGAGCATCCACCAAG |
| Hk1 | GCCACGCCTCGGTGCCATCTT | GGTCTTGTGGAACCGCCGGG |
| Hprt | TGCCGAGGATTTGGAAAAAGTG | CACAGAGGGCCACAATGTGATG |
| Ikzf1 (Ikaros) | GGAGGCACAAGTCTGTTGAT | CATTTCACAGGCACGCCCATTCT |
| Ikzf3 (Aiolos) | GCCGAGATGGGAAGTGAGAG | CCGGGATTGTAGTTGGCATC |
| Myc | CGAAACTCTGGTGCATAAACTG | GAACCGTTCTCCTTAGCTCTCA |
| Rag1 | GGGGAGTGGGGTTGAAAGTA | TCCTCCAATCCTGCCTCCTA |
| Slc2a1 (Glut1) | AGCCCTGCTACAGTGTAT | AGGTCTCGGGTCACATC |
| Txnip | CGAGTCAAAGCCGTCAGGAT | TTCATAGCGCAAGTAGTCCAAAGT |
